# *Thiovibrio frasassiensis* gen. nov., sp. nov., an autotrophic, elemental sulfur disproportionating bacterium isolated from sulfidic karst sediment, and proposal of Thiovibrionaceae fam. nov.

**DOI:** 10.1101/2022.10.27.514068

**Authors:** H.S. Aronson, C. Thomas, M.K. Bhattacharyya, S.R. Eckstein, S.R. Jensen, R.A. Barco, J.L. Macalady, J.P. Amend

## Abstract

A novel, autotrophic, mesophilic bacterium, strain RS19-109^T^, was isolated from sulfidic stream sediments in the Frasassi Caves, Italy. The cells of this strain grew chemolithoautotrophically under anaerobic conditions while disproportionating elemental sulfur (S^0^) and thiosulfate, but not sulfite with bicarbonate/CO_2_ as a carbon source. Autotrophic growth was also observed with molecular hydrogen as an electron donor, and S^0^, sulfate, thiosulfate, nitrate, and ferric iron as electron acceptors. Oxygen was not used as an electron acceptor and sulfide was not used as an electron donor. Weak growth was observed with sulfate as an electron acceptor and organic carbon as electron donors and carbon sources. The strain also showed weak growth by fermentation of tryptone. Strain RS19-109^T^ was found to be phylogenetically distinct based on 16S rRNA gene sequence similarity (89.2%) to its closest relative, *Desulfurivibrio alkaliphilus* AHT2^T^. The draft genome sequence for strain RS19-109^T^ had average nucleotide identity, average amino acid identity, and *in silico* DNA-DNA hybridization values of 72.2%, 63.0%, and 18.3%, respectively, compared with the genome sequence of *D. alkaliphilus* AHT2^T^. On the basis of its physiological and genomic properties, strain RS19-109^T^ is proposed as the type strain of a novel species of a novel genus, *Thiovibrio frasassiensis* gen. nov., sp. nov. A novel family, *Thiovibrionaceae* fam. nov., is proposed to accommodate *Thiovibrio* within the order Desulfobulbales.

## Introduction

Sulfur disproportionation, or the simultaneous oxidation and reduction of intermediate oxidation state sulfur compounds to form sulfate and sulfide, is a relatively understudied branch of the sulfur cycle compared to sulfate reduction and sulfide oxidation. Most microorganisms capable of sulfur disproportionation are able to use thiosulfate and sulfite, but the ability to disproportionate elemental sulfur (S^0^) is much less common (Slobodkin & Slobodkina, 2019). To date, 20 S^0^-disproportionating bacteria within the phyla Desulfobacterota, Proteobacteria, Firmicutes, and Nitrospirae have been isolated from freshwater sediments (Widdel & Pfennig, 1982; Lovley & Phillips, 1994; Janssen *et al*., 1996), marine sediments (Friedrich *et al*., 1996; Finster *et al*., 1998; Francis *et al*., 2000; Obraztsova *et al*., 2002), freshwater and soda lakes (Peduzzi *et al*., 2003; Sorokin *et al*., 2008; Poser *et al*., 2013), hydrothermal vents (Slobodkin *et al*., 2012, 2013; Slobodkina *et al*., 2016, 2017), acidic river sediments (Florentino *et al*., 2016), and hot springs (Kojima *et al*., 2016; Merkel *et al*., 2017; Umezawa *et al*., 2021). In this study, we report the isolation and characterization of a novel S^0^ disproportionating bacterium, strain RS19-109^T^, from sulfidic stream sediments in the karst at Frasassi, Italy. This is the first S^0^ disproportionating microorganism isolated from the terrestrial subsurface. Based on its physiological features and genomic sequence, this strain represents a novel genus and species for which we proposed the name *Thiovibrio frasassiensis*, which we propose to place within a novel family which we designate *Thiovibrionaceae*.

## Methods

### Enrichment and Isolation

Strain RS19-109^T^ was isolated from a sediment sample collected from a shallow sulfidic stream in the Frasassi Caves, Italy (43.4012 N, 12.9656 E) in July 2019. The sampling point was located at Ramo Sulfureo in Grotta del Fiume at 400 meters below surface (Macalady *et al*., 2006). Water temperature at the sampling site was 13ºC, and the pH of the water was 7.3. Sediment samples were collected using a syringe, transferred into serum bottles purged with nitrogen and sealed with butyl stoppers, and transported back to the laboratory at the Geological Observatory at Coldigioco, Italy.

For enrichment of sulfur disproportionators, a bicarbonate-buffered freshwater minimal medium was prepared under 80% N_2_-20% CO_2_. The medium was based on DSMZ 386 but acetate was omitted, amorphous ferric hydroxide was added as a sulfide scavenger, and sodium pyrosulfite was replaced with 10 g/L S^0^. The basal medium contained the following (per liter of deionized water): 0.20 g KH_2_PO_4_, 0.25 g NH_4_Cl, 1.0 g NaCl, 0.40 g MgCl_2_ x 6 H_2_O, 0.50 g KCl, 0.15 g CaCl_2_ x 2 H_2_O, 1 mL trace element solution SL-10 (see DSMZ medium 320), 1 mL selenite-tungstate solution (see DSMZ medium 385), and 1 mL DSMZ seven vitamin mixture solution (see DSMZ medium 503). Amorphous ferric hydroxide was prepared as described by Lovley and Phillips, 1986 and was added to the medium at a final concentration of 30 mM (Lovley & Phillips, 1986). The pH of the medium was adjusted to 7.0 using NaOH. Medium was dispensed into serum bottles containing 10 g/L S^0^ and was sterilized by heating for 2-3 hours in a boiling water bath on each of three successive days. After sterilization, sodium bicarbonate was added at a final concentration of 30 mM and sodium sulfide was added as a reductant (final concentration 0.50 μM). The complete medium was inoculated with Frasassi sediment (1% w/v).

After 3 weeks of incubation at 15ºC, cell densities increased from 10^4^ cells ml^-1^ to 10^11^ cells ml^-1^ and the medium color changed from rusty red to black, indicating a reduction of ferric iron. After 40 days, average ferrous iron concentrations increased to 1.825 mM and average sulfate concentrations increased to over 21.37 mM (Fig. 1). Measurements of ferrous iron using the ferrozine assay were used as partial proxies for sulfide concentration (Stookey, 1970) and sulfate concentrations were measured using a turbidimetric sulfate assay (Kolmert *et al*., 2000). The culture was purified by dilution-to-extinction. Purity of the resulting isolate was confirmed by microscopy and Sanger sequencing of the 16S rRNA gene, and the resulting isolate was designated as strain RS19-109^T^. Cells of strain RS19-109^T^ were vibrio-shaped bacteria.

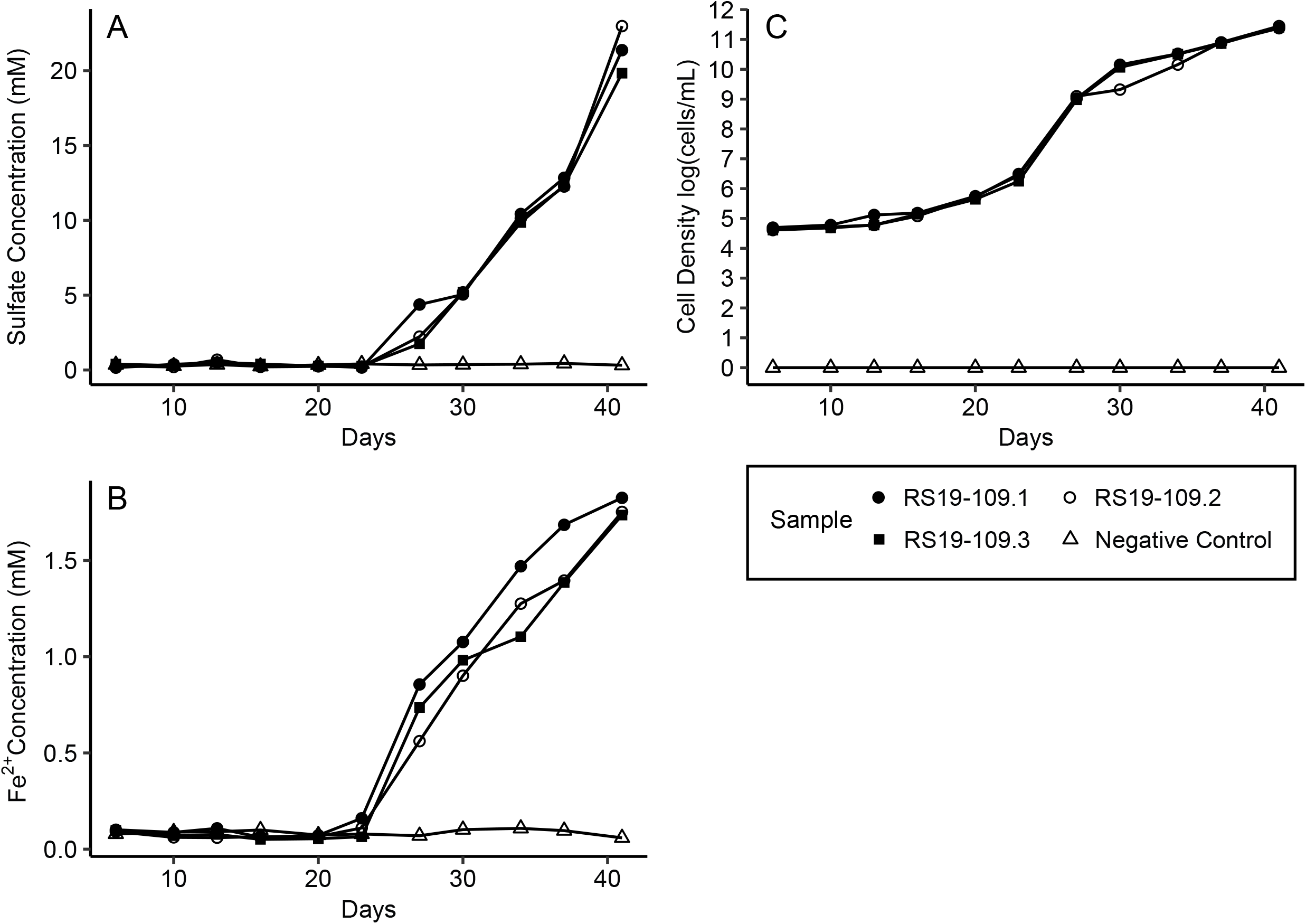

### Light and electron microscopy

Growth of strain RS19-109^T^ was determined by direct counts using fluorescence microscopy. 1 mL of culture was fixed in glutaraldehyde (2.5% final concentration) and stained with 125 μL of 10X SYBR Green I. Cultures with ferric iron were treated with an oxalate solution (28 g ammonium oxalate and 15 g anhydrous oxalic acid per liter) prior to staining to remove iron particles (Lovley & Phillips, 1988). 400 μL of oxalate solution was added to 1 mL of culture and was incubated for 15 minutes at room temperature, neutralized with NaOH, and stained with 125 μL of 10X SYBR Green I. Samples were filtered through black polycarbonate filters with a 0.22 μm pore size and cells were counted under a fluorescence microscope.

Scanning electron microscopy was used to obtain images of strain RS19-109^T^ cells (Fig. 2). Cells were fixed in 2% glutaraldehyde for 1 hour at room temperature, centrifuged for 2 minutes at 10,000 rpm to pellet the minerals, and the supernatant was filtered onto a 0.1 μm Supor filter. Samples were dehydrated in an ethanol series (30, 50, 70, 80, 90, 95, and 100%), dried using a Tousimis 815 critical point dryer, and sputter-coated for 40 seconds with Pt/Pd using a Cressington 108 manual sputter coater. Cells were observed with a Nova NanoSEM 450 field emission scanning electron microscope at an accelerating voltage of 5.0 kV.

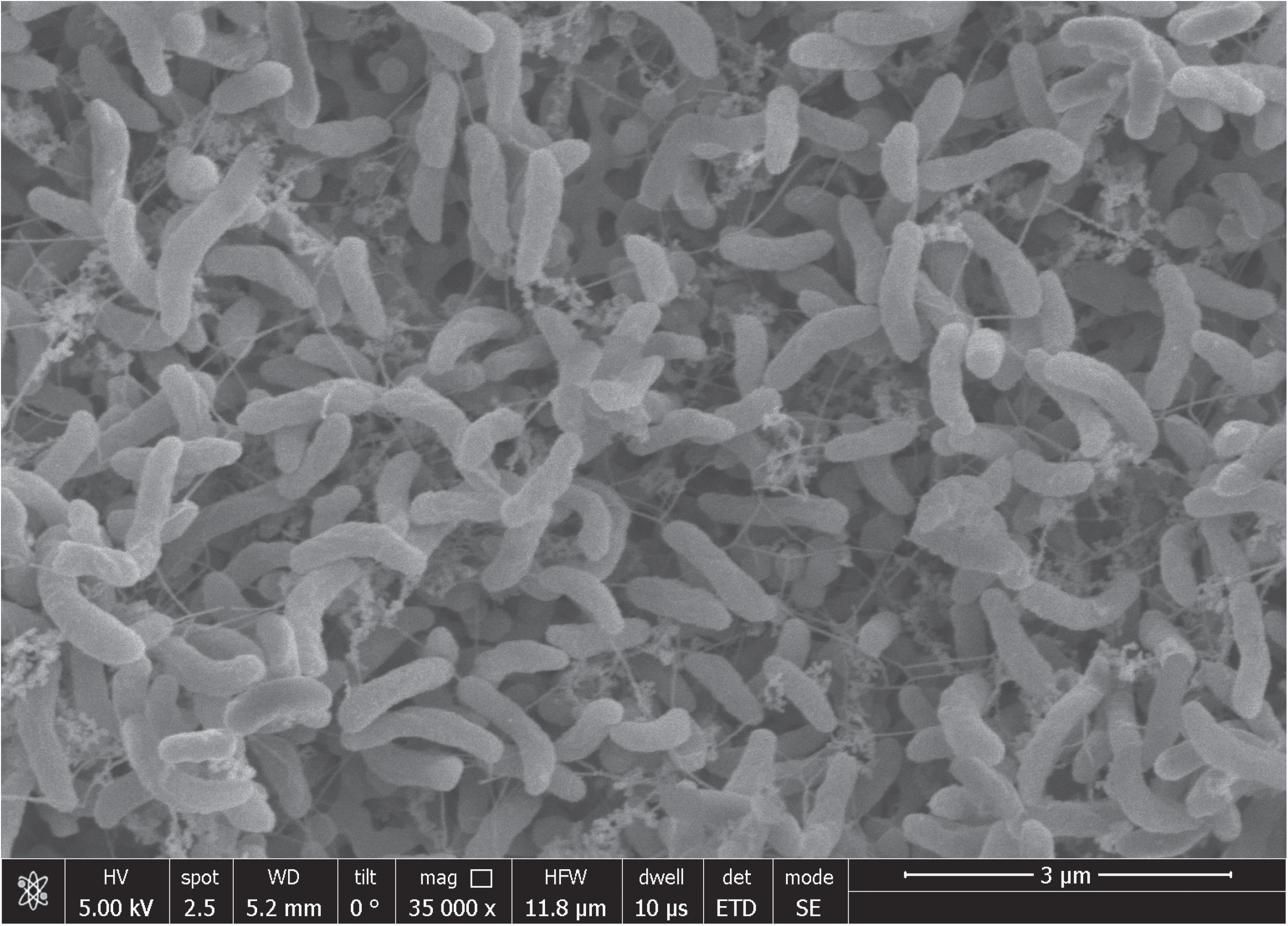

### Temperature, pH, and salt tolerance

Growth experiments to evaluate tolerance for temperature, pH, and salinity were performed in triplicate in serum bottles with the same basal medium used for pure culture isolation. The temperature range for growth was evaluated at 4ºC-50ºC at 5ºC intervals at pH 7.0. The temperature optimum was 30ºC. No growth was detected above 36ºC after incubation for 4 weeks. The pH range for growth was evaluated at 30ºC over a range of pH 5.0 to 8.5 at intervals of 0.5 pH unit. The culture medium was supplemented with the following buffers at final concentrations of 25 mM: MES (2-(*N*-morpholino)ethanesulfonic acid; pH 5.0-6.5 and HEPES (4-(2-hydroxyethyle)-1-piperazineethanesulfonic acid; 7.0-8.5. The pH optimum was 7.0 and no growth was observed below pH 5.5 or above pH 7.5. Salinity tolerance of strain RS19-09^T^ was evaluated at 30ºC using NaCl-free basal growth medium supplemented with NaCl (0, 0.1, 0.25, 0.5, 0.75, 1, 1.2, 1.5, 1.7, 2.0%). The salinity optimum was 0-1.7% NaCl. No growth was observed above 1.7% NaCl.

### Physiology and chemotaxonomy

All phenotypic tests were performed in triplicate in serum bottles. Chemolithoautotrophic growth on S^0^ as a substrate for disproportionation was tested in the basal medium with 10 g/L S^0^ added as the electron donor and acceptor and bicarbonate/CO_2_ added as the carbon source and amorphous ferric oxyhydroxide as a sulfide scavenger. Disproportionation of thiosulfate and sulfite was tested in the basal medium with S^0^ replaced with 20 mM thiosulfate or sulfite. Strain RS19-109^T^ was able to disproportionate S^0^ and thiosulfate, but not sulfite. Growth was accompanied by production of sulfate and conversion of amorphous ferric hydroxide to a black precipitate, which was likely composed of both acid-volatile sulfides (H_2_S and FeS) and chromium (II)-reducible sulfides (S^0^ and pyrite, FeS_2_) (Thamdrup *et al*., 1993). S^0^, which was present at 10 g/L, would likely cause a high background concentration of sulfide in the CRS fraction, so the ferrozine assay was used to measure HCl-extractable Fe^2+^ as a proxy for acid-volatile sulfide concentration. Total sulfide concentration can be calculated as the sum of AVS and (1/2 x CRS) because only half of the sulfur in pyrite exists at the oxidation state of sulfide (−2). By the end of the experiment, all of the red amorphous ferric oxyhydroxide (30 mM) turned black and the final average measured Fe^2+^ concentration was 1.825 mM, suggesting that 56.35 mM of CRS was produced (although this would also include an undetermined amount of S^0^).

Utilization of electron donors with sulfate (20 mM) as the electron acceptor was tested using bicarbonate-free basal medium with the addition of 10 mM of each of the following substrates to the medium: acetate, citrate, ethanol, formate, fumarate, glucose, lactate, pyruvate, succinate, tryptone, and yeast extract (1%); the test was conducted with and without the addition 0.001% yeast extract. The use of carbon source as an electron donor for sulfate reduction or as a substrate for fermentation was determined by measurements of sulfide with the Cline methylene blue assay (Cline, 1969). Strain RS19-109^T^ showed weak growth as a sulfate reducer using acetate, ethanol, formate, fumarate, glucose, pyruvate and yeast extract (0.1%). It did not reduce sulfate with tryptone, but instead fermented tryptone which was accompanied by weak growth without the production of sulfide. No growth was observed with citrate, lactate, or succinate.

Electron acceptor tests were performed with 20 mM of the following compounds in the basal medium with N_2_/CO_2_ replaced with 80% H_2_-20% CO_2_. Strain RS19-109^T^ used H_2_ as an electron donor with S^0^, sulfate, thiosulfate, nitrate, and amorphous ferric hydroxide as electron acceptors. It was also able to reduce ferric iron with acetate (10 mM). Sulfite was not used as an electron acceptor. Acetate was not used as an electron donor for nitrate reduction, with or without the addition of 0.001% yeast extract. Aerobic respiration was tested with S^0^ and sulfide as electron donors in the basal medium. Oxygen was added to the headspace by placing a needle fitted with a syringe filter through the butyl rubber stopper; strain RS19-109^T^ was incapable of aerobic respiration.

### 16S rRNA phylogeny and genome features

Genomic DNA was extracted from strain RS19-109^T^ using the Qiagen DNeasy UltraClean Microbial Kit and was used for 16S rRNA gene sequencing and whole genome sequencing. The 16S rRNA gene was amplified using 27F/1492R primers and was sequenced by GeneWiz (South Plainfield, NJ). 16S rRNA gene sequences from close isolated relatives identified using BLAST from the NCBI ref_seq_rna database were aligned using MUSCLE v3.8.31 (Edgar, 2004). Gaps were removed and sequences were trimmed using trimAl v1.2 (Capella-Gutiérrez *et al*., 2009). A phylogenetic tree was constructed using the aligned and trimmed 16S rRNA gene sequences using IQ-TREE with a maximum-likelihood method and the generalized time-reversible model of nucleotide evolution with 1000 bootstrap replications (Fig. 3) (Nguyen *et al*., 2015). To obtain 16S rRNA gene sequence identities, sequences were aligned using the SILVA SINA aligner v.1.2.11 (Pruesse *et al*., 2012; Quast *et al*., 2012), gaps were removed, and identities were calculated in Geneious v6.1.8 (https://www.geneious.com).

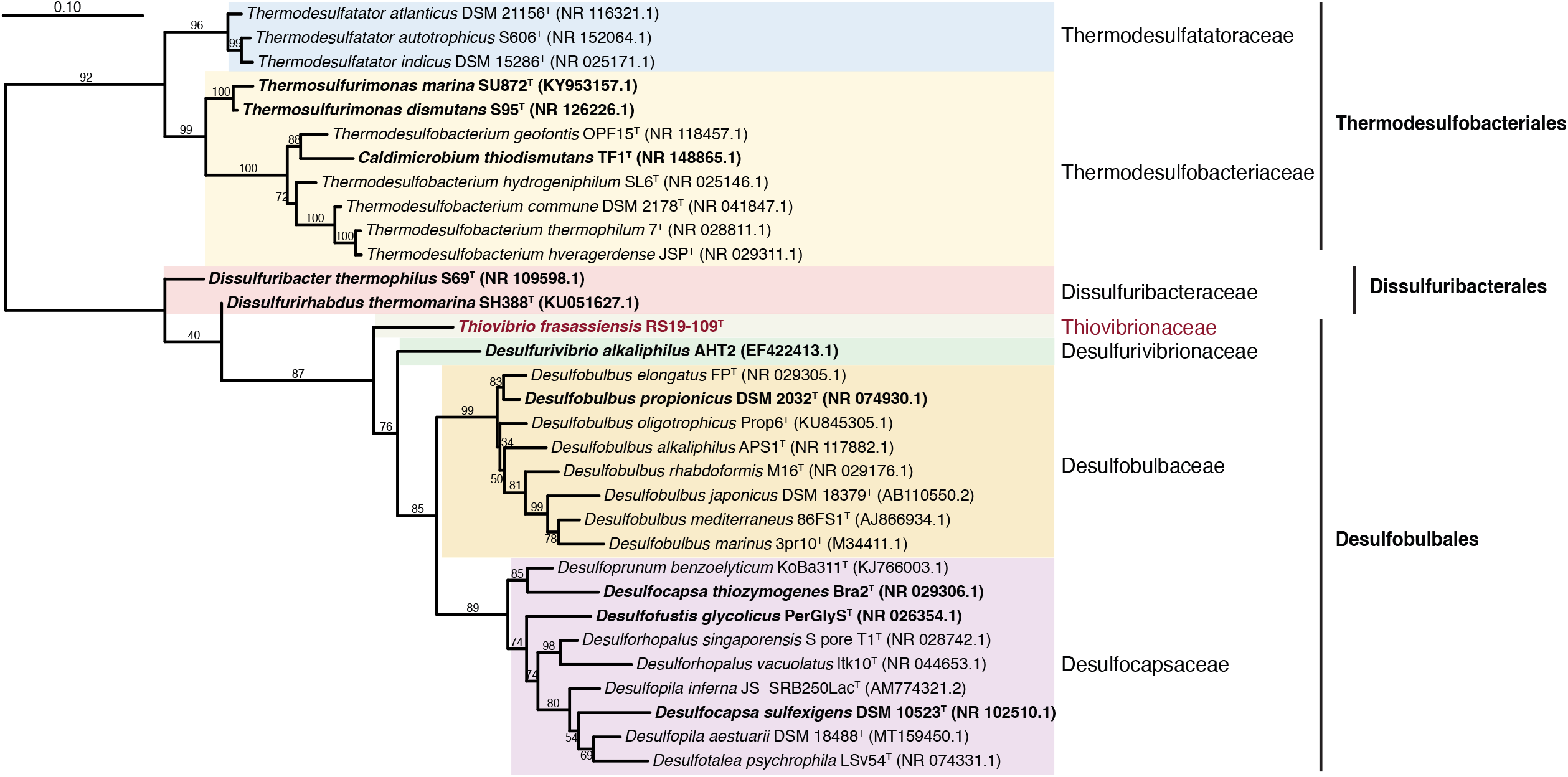

The genome of strain RS19-109^T^ was sequenced by Novogene (Sacramento, CA) on an Illumina NovoSeq platform yielding 7,079,420 paired-end reads. The sequence quality of the reads was determined with FastQC v0.11.8 (Simons, 2010). The read length was 150 bp. Error-corrected reads were assembled with SPAdes v3.15.4 (Prjibelski *et al*., 2020). Genome quality was determined using CheckM (Parks *et al*., 2015) and was 98.13% complete and 0.6% contaminated. The draft genome sequence comprised of 2,841,614 bp within 3 contigs. The G+C content of the genomic DNA was 56.25, the N50 was 2,800,286, and the sequence depth of coverage was 374X. 2621 protein-coding sequences and 48 tRNA-encoding genes were predicted. The draft genome of strain RS19-109^T^ contained one copy each of 5S, 16S, and 23S rRNA genes.

Closely-related strains were identified using GToTree (Lee, 2019). The “gtt-get-accessions-from-GTDB” command was used to collect representative genomes from the order Desulfobulbales that were present in the Genome Taxonomy Database (Parks *et al*., 2022). Pairwise average nucleotide identity (ANI), average amino acid identity (AAI), and digital DNA-DNA hybridization (dDDH) values were calculated for all genomes in the Desulfobulbales using the IMG ANIcalculator (Varghese *et al*., 2015), EzAAI (Kim *et al*., 2021), and the Genome-to-Genome Distance Calculator v3.0 (Meier-Kolthoff *et al*., 2013, 2022), respectively. Genomes from isolated species, and genomes from the closest uncultured relatives of strain RS19-109^T^ were used to construct a phylogenomic tree (Fig. 4). GTDB-tk (v2.1.1) was used to align 120 conserved single-copy marker gene sequences (bac120) from the genomes (Parks *et al*., 2018, 2022; Chaumeil *et al*., 2020). IQ-TREE was used to construct the tree from the aligned sequences using the methods described above (Nguyen *et al*., 2015).

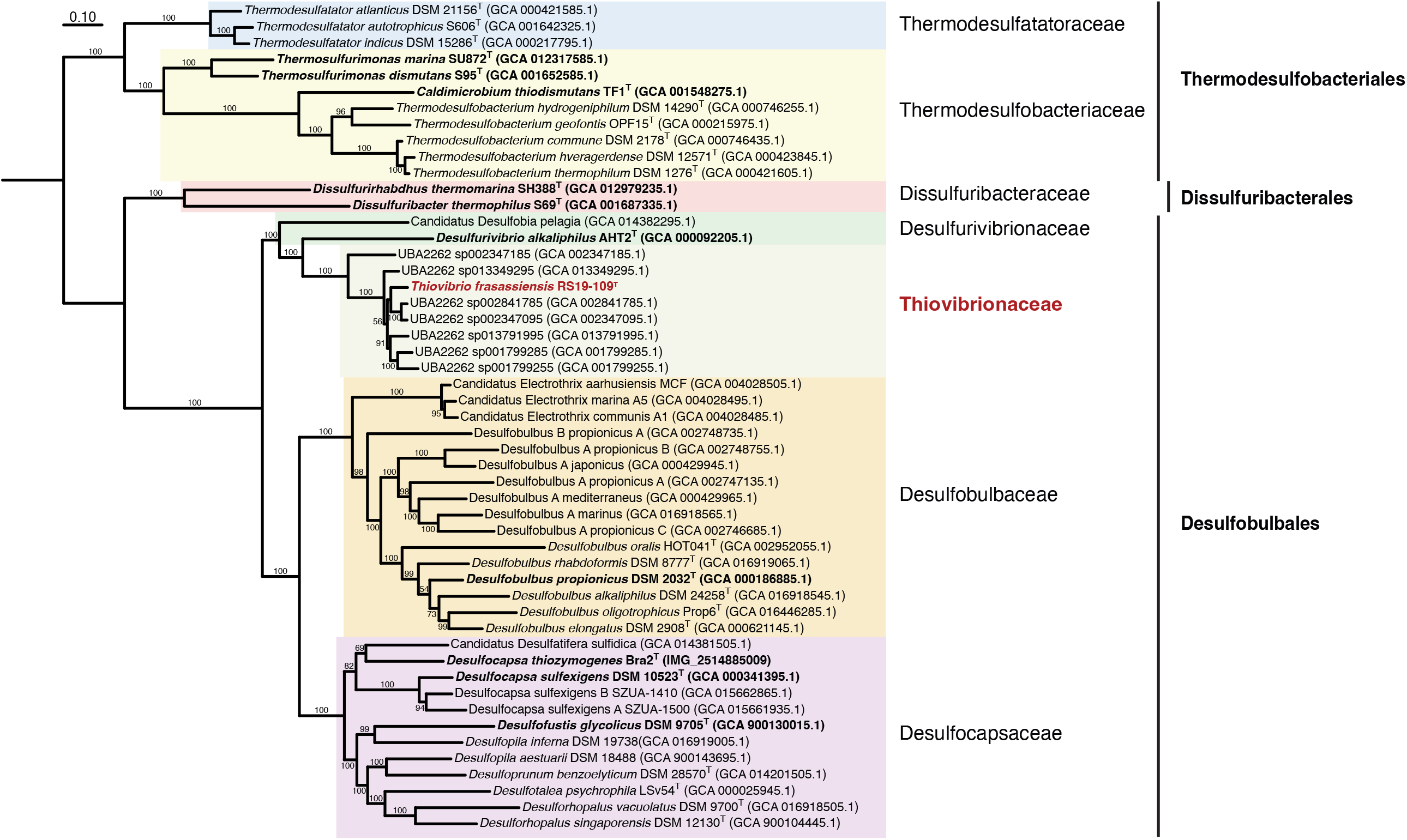

The closest recognized species to strain RS19-109^T^ based on 16S rRNA gene sequence similarity was *Desulfurivibrio alkaliphilus* AHT2^T^ (89.2%) (Sorokin *et al*., 2008), which is far below the threshold of 98.65% for differentiating two species (Kim *et al*., 2014; Konstantinidis *et al*., 2017) and the 94.5% sequence similarity for designating the rank of genus (Yarza *et al*., 2014). Using the median value for the rank of family designated by Yarza and colleagues (<92.25%) and the range for novel families designated by Konstantinidis and colleagues (89-92%), we propose that strain RS19-109^T^ belongs to a novel species, genus, and family. In a phylogenomic tree, strain RS19-109^T^ and its closest uncultured relatives formed a clade that clustered most closely with the family Desulfurivibrionaceae, outside of the families Desulfobulbaceae and Desulfocapsaceae (Fig. 3) (Waite *et al*., 2020). For genomic discreteness, the ANI value (72.2%) and dDDH value (18.3%) of strain RS19-109^T^ fall well below the species-level ANI cutoff and the dDDH cutoff for relatedness to its closest isolated relative (<95% and <70%, respectively) (Table 2, Supp. Files 1 and 2) (Konstantinidis *et al*., 2017). The proposed AAI thresholds for designating same genera and families are 65-95% and 45-65%, respectively (Konstantinidis *et al*., 2017). When compared to the type genera of the families within the Desulfobulbales designated by Waite et al. 2020, strain RS19-109^T^ has AAI values of 58.6 to 63%, which would classify it as a novel genus within the family Desulfurivibrionaceae. However, when these type genera are compared to each other, *D. alkaliphilus* AHT2^T^, *D. propionicus* 1pr3^T^, and *D. thiozymogenes* Bra2^T^ all had AAI values to each other and to RS19-109^T^ of at least 58%, putting them all well above the 45% AAI family level cutoff that would designate them as novel families (Table 2; Konstantinidis *et al*., 2017). On the basis of its phylogenomic placement and 16S rRNA gene sequence similarity to close relatives, we propose that strain RS19-109^T^ is placed within a novel family in the order Desulfobulbales, which we designate Thiovibrionaceae.

**Table 1.**
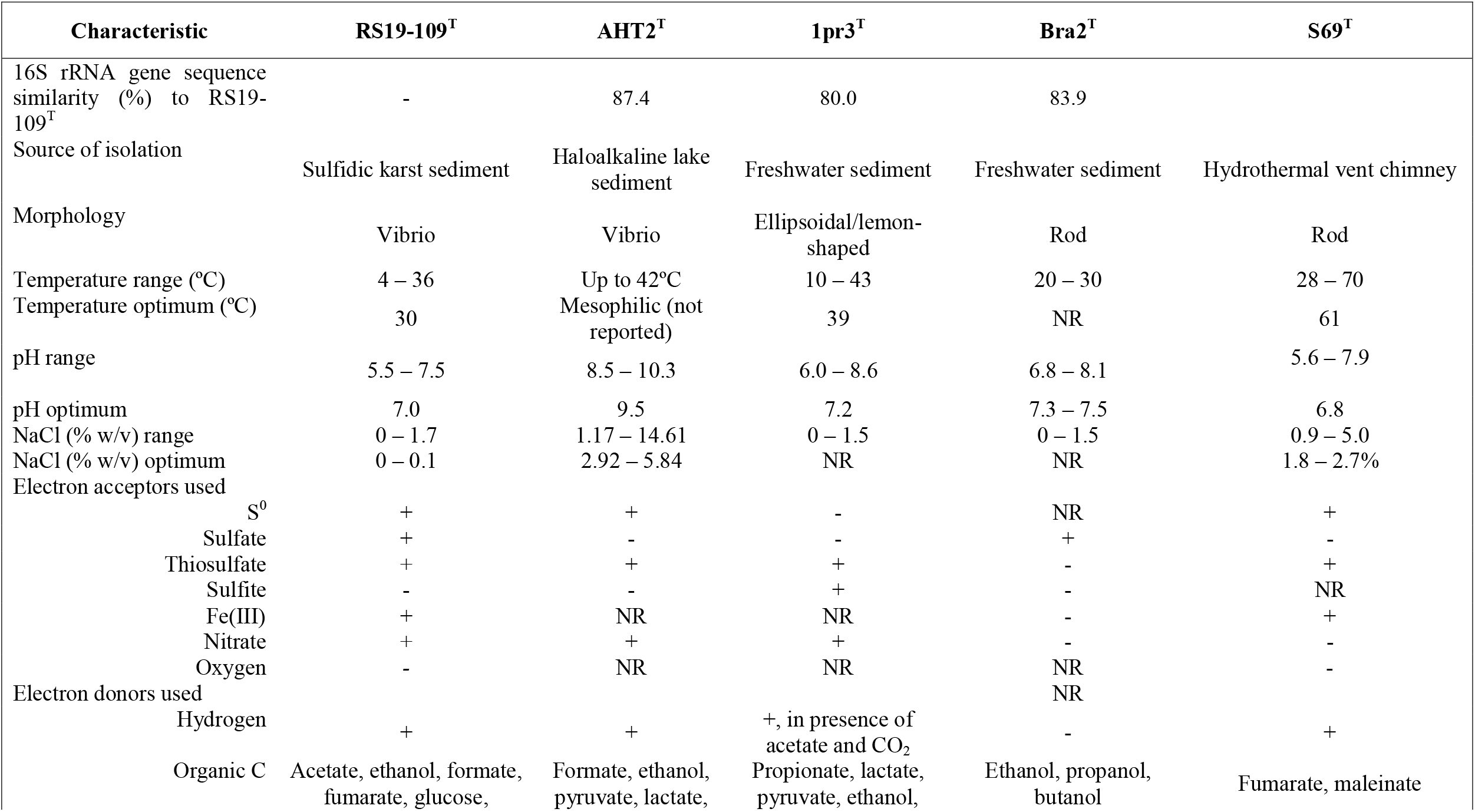
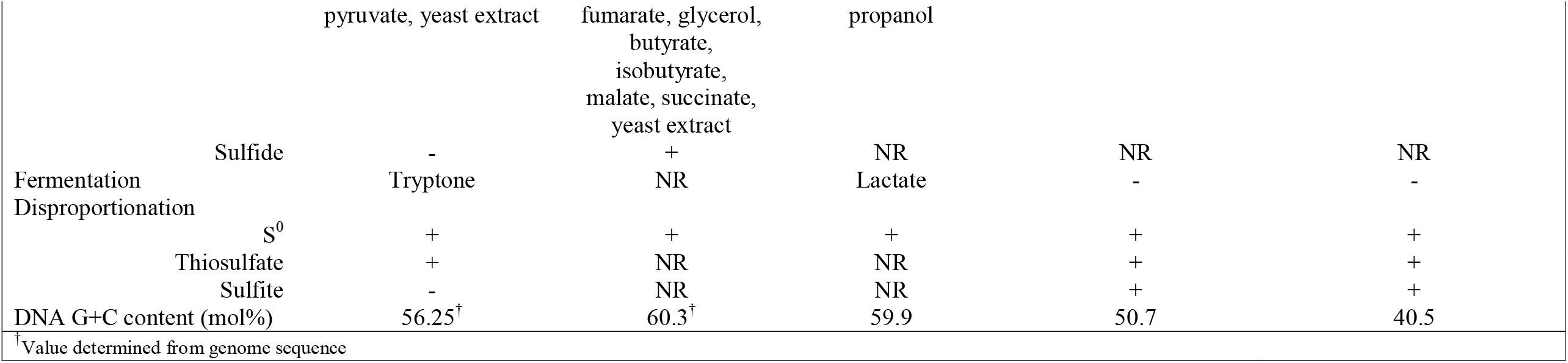
Characteristics that distinguish strain RS19-109^T^ from representatives of closely related species. Strains: RS19-109^T^ (data from this study); *Desulfurivibrio alkaliphilus* AHT2^T^ = DSM 19089 (data from Sorokin et al. 2008); *Desulfobulbus propionicus* 1pr3 = DSM 2032 (data from Widdel et al. 1981 and Lovley and Phillips 1994); *Desulfocapsa thiozymogenes* Bra2=DSM 7269 (data from Janssen et al. 1996); *Dissulfuribacter thermophilus* S69^T^=DSM 25762 (data from Slobodkin et al. 2013). NR = Not reported.

**Table 2.**
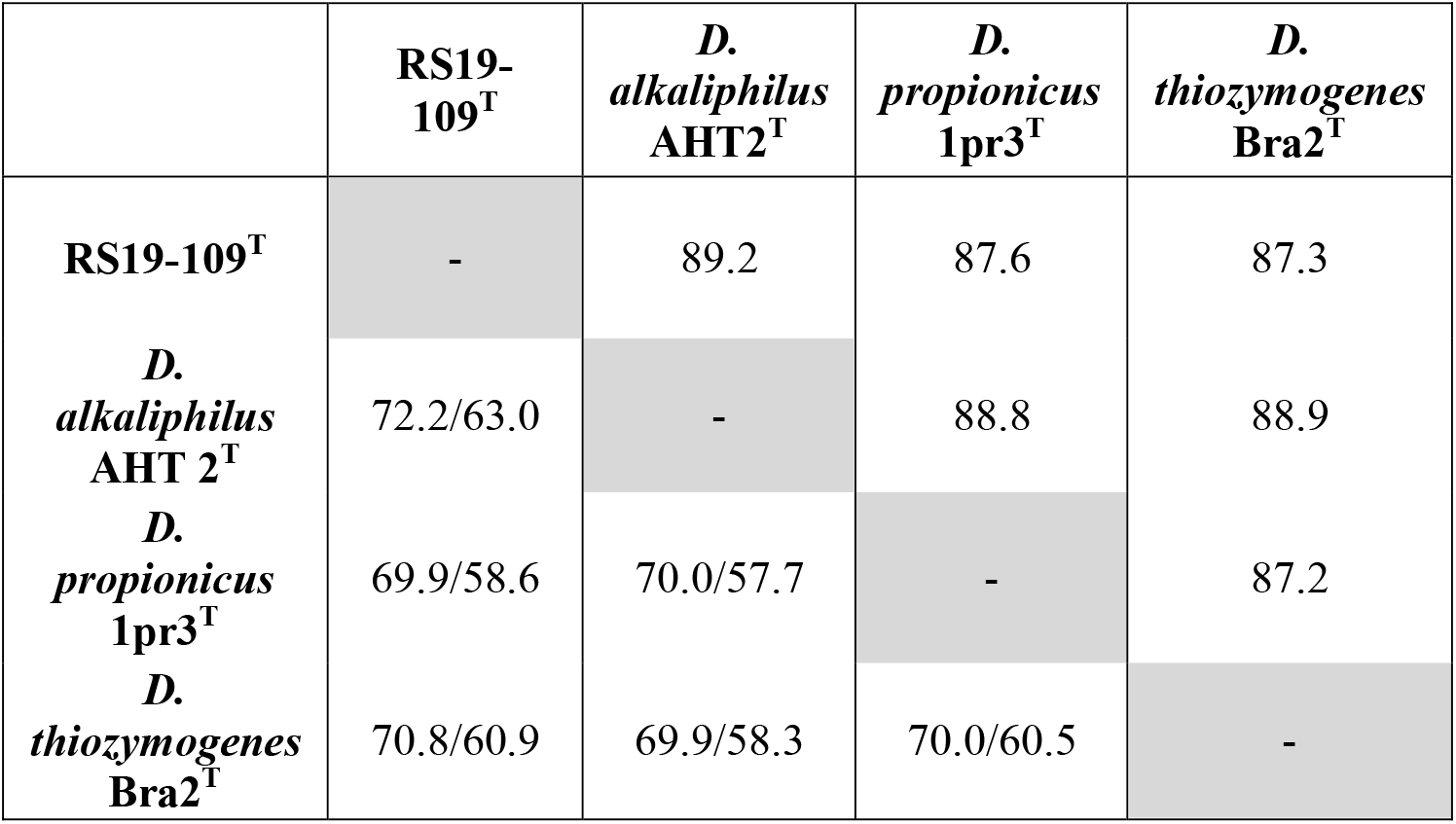
Pairwise comparisons of 16S rRNA gene sequence identities (%, values to the right of the gray boxes) and ANI/AAI (%, values to the left of the gray boxes) between strain RS19-109^T^ and type genera of the families Desulfurivibrionaceae, Desulfobulbaceae, and Desulfocapsaceae.

Annotation of the RS19-109^T^ draft genome was performed using MetaSanity (Neely *et al*., 2020) and METABOLIC (Supp. File 3) (Zhou *et al*., 2019). The genome of RS19-109^T^ contained genes encoding enzymes involved in carbon fixation via the Wood-Ljungdahl pathway (acetyl-CoA synthase *cdhDE/acs*; carbon monoxide dehydrogenase *cooS*). Strain RS19-109^T^ also has the genetic potential for acetogenesis (acetyl coA synthetase *acdA*; acetate kinase *ack*; phosphate acetyl transferase *pta*), formate oxidation (formate dehydrogenase *fdoH*), and pyruvate oxidation (pyruvate ferredoxin oxidoreductase *porA*). The genome contained genes encoding nitrogenases (*nifDKH*) and a dissimilatory nitrate reduction pathway (nitrate reductase *napAB;* nitrate reductase *nrfAH*).

Several genes encoding enzymes in sulfur metabolism were present in the genome of RS19-109^T^, including a complete dissimilatory sulfate reduction pathway (sulfate adenylyltransferase *sat*; adenylylsulfate reductase *aprAB*; dissimilatory sulfite reductase *dsrAB;* other dissimilatory sulfite reductase subunit genes *dsrCDJKMOPT*), sulfur oxidation (sulfur dioxygenase *sor*), and thiosulfate disproportionation (thiosulfate reductase/polysulfide reductase *phsA*). The genome also contained genes for arsenate reduction (arsenate reductase *arsC2*) and iron/manganese reduction (decaheme-associated outer membrane protein *mtrB*). Genes for hydrogen oxidation (Ni-Fe hydrogenase *nife*-group 1), cytochrome bd ubiquinol oxidase (*cydAB*), NADH-quinone oxidoreductase Complex I (nuoAB), and F-type H+ transporting ATP-ase (atpAD) were also present. The genes and pathways present in the genome are consistent with the ability of strain RS19-109^T^ to reduce sulfate with hydrogen, disproportionate thiosulfate, reduce iron, reduce nitrate, and utilize bicarbonate/CO_2_ as a carbon source. To date, genomic markers for S^0^ disproportionation have not been identified (Slobodkin & Slobodkina, 2019; Ward *et al*., 2021), and we were thus unable to ascribe any genes or pathways to the ability of RS19-109^T^ to disproportionate S^0^.

### Description of *Thiovibrio* gen. nov. *non Thiovibrio* (Janke, 1924*) non Thiovibrio* (Emoto & Hirose, 1942)

*Thiovibrio* (Thi.o.vi’bri.o.) Gr. Neut. n. *theion*, sulfur; N.L. masc. n. *vibrio*, a vibrio; N.L. masc. n. *Thiovibrio*, a sulfur vibrio.

Cells are vibrio shaped. Anaerobic, mesophilic, and neutrophilic. Chemolithoautotrophic growth by disproportionation of S^0^ to sulfide and sulfate. Member of the phylum Desulfobacterota. The type species is *Thiovibrio frasassiensis*.

### Description of *Thiovibrio frasassiensis*, sp. nov

*Thiovibrio frasassiensis* (fra.sa.si.en.sis. N.L. fem. adj. *frasassiensis* pertaining to Frasassi, the name of the caves from which the type strain was isolated).

Shows the following properties in addition to those given in the genus description. Cells are vibrio shaped and occur singly, 0.6 μm in diameter and 2.3 μm in length. The temperature range for growth is 4-36ºC, with an optimum at 30ºC. The pH range for growth is 5.5-7.5, with an optimum at pH 7.0. Grows in 0-1.7% (w/v) NaCl with an optimum at 0-0.1% (w/v). S^0^ and thiosulfate are disproportionated to form sulfate and sulfide with bicarbonate/CO_2_ as a carbon source. Growth is enhanced in the presence of amorphous ferric hydroxide. In the presence of 80% H_2_-20% N_2,_ S^0^, sulfate, thiosulfate, nitrate, and amorphous ferric hydroxide serve as electron acceptors. Weak anaerobic heterotrophic growth is observed on acetate, ethanol, formate, fumarate, glucose, pyruvate, and yeast extract with sulfate as the electron acceptor. Fermentative growth is observed on tryptone. Does not oxidize or ferment citrate, lactate, or succinate. Grows by the oxidation of H_2_ with S^0^, sulfate, thiosulfate, nitrate, and ferric iron. Does not grow by oxidizing sulfide or S^0^ with oxygen and does not use sulfite as an electron acceptor.

The type strain is RS19-109^T^, isolated from sulfidic stream sediment from the karst at Frasassi, Italy. The G+C content of the genomic DNA of the type strain is 56.25 mol% (determined from the genome sequence).

### Description of *Thiovibrionaceae* fam. nov

*Thiovibrionaceae* (Thi.o.vi’bri.o.na.ce’ae. *Thiovibrio* type genus of the family; -aceae ending to denote family) Encompasses chemolithoautotrophic bacteria. Based on 16S rRNA gene sequence and whole genome analysis the strain is phylogenetically affiliated with the order Desulfobulbales.

## Acknowledgements

The authors thank A. Montanari for providing logistical support and the use of facilities and laboratory space at the Osservatorio Geologico di Coldigioco, Apiro, Italy. Thanks to members of the Gruppo Speleologico C.A.I. di Fabriano for technical assistance during field campaigns, and to D. Jones, Z. Havlena, and M. Best for assistance in the field. H.S.A. was supported by an NSF Graduate Research Fellowship grant DGE-1842487, the Lewis and Clark Fund for Exploration and Field Research in Astrobiology, the National Cave and Karst Institute NCKRI Scholar Fellowship, the Josephine de Karman Fellowship, and the Cave Conservancy Foundation Graduate Fellowship in Karst Studies. MB was supported by the USC Summer Undergraduate Research Fund. Electron micrographs were acquired at the Core Center of Excellence in Nano Imaging at the University of Southern California with technical assistance from Daniel Goodelman.

## References

Capella-Gutiérrez S, Silla-Martínez JM, Gabaldón T (2009) trimAl : a tool for automated alignment trimming in large-scale phylogenetic analyses 25, 1972–1973.

Chaumeil P-A, Mussig AJ, Hugenholtz P, Parks DH (2020) GTDB-Tk: a toolkit to classify genomes with the Genome Taxonomy Database. Bioinformatics 36, 1925–1927.

Cline JD (1969) Spectrophotometric Determination of Hydrogen Sulfide in Natural Waters. Limnology and Oceanography 14, 454–458.

Edgar RC (2004) MUSCLE: Multiple sequence alignment with high accuracy and high throughput. Nucleic Acids Research 32, 1792–1797.

Emoto Y, Hirose H (1942) Studien über die Thermalflora von Japan. XIX. Thermal Bakterien und Algen aus thermal Quellen von Hatimantai und Yakeyama. Bot. Mag. Tokyo 332–343.

Finster K, Liesack W, Thamdrup B (1998) Elemental sulfur and thiosulfate disproportionation by Desulfocapsa sulfoexigens sp. nov., a new anaerobic bacterium isolated from marine surface sediment. Applied and environmental microbiology 64, 119–125.

Florentino AP, Brienza C, Stams AJM, Sánchez-Andrea I (2016) Desulfurella amilsii sp. nov., a novel acidotolerant sulfur-respiring bacterium isolated from acidic river sediments. International Journal of Systematic and Evolutionary Microbiology 66, 1249–1253.

Francis CA, Obraztsova AY, Tebo BM (2000) Dissimilatory Metal Reduction by the Facultative Anaerobe Pantoea agglomerans SP1. Applied and Environmental Microbiology 66, 543–548.

Friedrich M, Springer N, Ludwig W, Schink B (1996) Phylogenetic Positions of Desulfofustis glycolicus gen. nov., sp. nov. and Syntrophobotulus glycolicus gen. nov., sp. nov., Two New Strict Anaerobes Growing with Glycolic Acid. International Journal of Systematic Bacteriology 46, 1065–1069.

Janke A (1924) Allgemeine technische Mikrobiologie. 1. Teil. Die Mikroorganismen. Steinkopff.

Janssen PH, Schuhmann A, Bak F, Liesack W (1996) Disproportionation of inorganic sulfur compounds by the sulfate-reducing bacterium Desulfocapsa thiozymogenes gen, nov., sp. nov. Archives of Microbiology 166, 184–192.

Kim D, Park S, Chun J (2021) Introducing EzAAI: a pipeline for high throughput calculations of prokaryotic average amino acid identity. Journal of Microbiology 59, 476–480.

Kim M, Oh H-S, Park S-C, Chun J (2014) Towards a taxonomic coherence between average nucleotide identity and 16S rRNA gene sequence similarity for species demarcation of prokaryotes. International Journal of Systematic and Evolutionary Microbiology 64, 346–351.

Kojima H, Umezawa K, Fukui M (2016) Caldimicrobium thiodismutans sp. nov., a sulfur-disproportionating bacterium isolated from a hot spring, and emended description of the genus Caldimicrobium. International Journal of Systematic and Evolutionary Microbiology 66, 1828–1831.

Kolmert Å, Wikström P, Hallberg KB (2000) A fast and simple turbidimetric method for the determination of sulfate in sulfate-reducing bacterial cultures. Journal of Microbiological Methods 41, 179–184.

Konstantinidis KT, Rosselló-Móra R, Amann R (2017) Uncultivated microbes in need of their own taxonomy. The ISME Journal 11, 2399–2406.

Lee MD (2019) GToTree: a user-friendly workflow for phylogenomics. Bioinformatics 35, 4162–4164.

Lovley DR, Phillips EJP (1986) Organic Matter Mineralization with Reduction of Ferric Iron in Anaerobic Sediments. Applied and Environmental Microbiology 51, 683–689.

Lovley DR, Phillips EJP (1988) Novel Mode of Microbial Energy Metabolism: Organic Carbon Oxidation Coupled to Dissimilatory Reduction of Iron or Manganese. Appl. Envir. Microbiol. 54, 1472–1480.

Lovley DR, Phillips EJP (1994) Novel processes for anaerobic sulfate production from elemental sulfur by sulfate-reducing bacteria. Applied and Environmental Microbiology 60, 2394–2399.

Macalady JL, Lyon EH, Koffman B, Albertson LK, Meyer K, Galdenzi S, Mariani S (2006) Dominant microbial populations in limestone-corroding stream biofilms, Frasassi cave system, Italy. Applied and Environmental Microbiology 72, 5596–5609.

Meier-Kolthoff JP, Auch AF, Klenk H-P, Göker M (2013) Genome sequence-based species delimitation with confidence intervals and improved distance functions. BMC Bioinformatics 14, 60.

Meier-Kolthoff JP, Carbasse JS, Peinado-Olarte RL, Göker M (2022) TYGS and LPSN: a database tandem for fast and reliable genome-based classification and nomenclature of prokaryotes. Nucleic Acids Research 50, D801–D807.

Merkel AY, Pimenov NV, Rusanov II, Slobodkin AI, Slobodkina GB, Tarnovetckii IY, Frolov EN, Dubin AV, Perevalova AA, Bonch-Osmolovskaya EA (2017) Microbial diversity and autotrophic activity in Kamchatka hot springs. Extremophiles 21, 307–317.

Neely CJ, Graham ED, Tully BJ (2020) MetaSanity: an integrated microbial genome evaluation and annotation pipeline. Bioinformatics 36, 4341–4344.

Nguyen LT, Schmidt HA, Von Haeseler A, Minh BQ (2015) IQ-TREE: A Fast and Effective Stochastic Algorithm for Estimating Maximum-Likelihood Phylogenies. Molecular Biology and Evolution 32, 268–274.

Obraztsova AY, Francis CA, Tebo BM (2002) Sulfur Disproportionation by the Facultative Anaerobe Pantoea agglomerans SP1 as a Mechanism for Chromium(VI) Reduction. Geomicrobiology Journal 19, 121–132.

Parks DH, Chuvochina M, Rinke C, Mussig AJ, Chaumeil P-A, Hugenholtz P (2022) GTDB: an ongoing census of bacterial and archaeal diversity through a phylogenetically consistent, rank normalized and complete genome-based taxonomy. Nucleic Acids Research 50, D785–D794.

Parks DH, Chuvochina M, Waite DW, Rinke C, Skarshewski A, Chaumeil P-A, Hugenholtz P (2018) A standardized bacterial taxonomy based on genome phylogeny substantially revises the tree of life. Nature Biotechnology.

Parks DH, Imelfort M, Skennerton CT, Hugenholtz P, Tyson GW (2015) CheckM: Assessing the quality of microbial genomes recovered from isolates, single cells, and metagenomes. Genome Research 25, 1043–1055.

Peduzzi S, Tonolla M, Hahn D (2003) Isolation and characterization of aggregate-forming sulfate-reducing and purple sulfur bacteria from the chemocline of meromictic Lake Cadagno, Switzerland. FEMS Microbiology Ecology 45, 29–37.

Poser A, Lohmayer R, Vogt C, Knoeller K, Planer-Friedrich B, Sorokin D, Richnow HH, Finster K (2013) Disproportionation of elemental sulfur by haloalkaliphilic bacteria from soda lakes. Extremophiles 17, 1003–1012.

Prjibelski A, Antipov D, Meleshko D, Lapidus A, Korobeynikov A (2020) Using SPAdes De Novo Assembler. Current Protocols in Bioinformatics 70, e102.

Pruesse E, Peplies J, Glöckner FO (2012) SINA: Accurate high-throughput multiple sequence alignment of ribosomal RNA genes. Bioinformatics 28, 1823–1829.

Quast C, Pruesse E, Yilmaz P, Gerken J, Schweer T, Yarza P, Peplies J, Glöckner FO (2012) The SILVA ribosomal RNA gene database project: improved data processing and web-based tools. Nucleic Acids Research 41, D590–D596.

Simons A (2010) A quality control tool for high throughput sequence data. A quality control tool for high throughput sequence data.

Slobodkin AI, Reysenbach AL, Slobodkina GB, Baslerov RV, Kostrikina NA, Wagner ID, Bonch-Osmolovskaya EA (2012) Thermosulfurimonas dismutans gen. nov., sp. nov., an extremely thermophilic sulfur-disproportionating bacterium from a deep-sea hydrothermal vent. International Journal of Systematic and Evolutionary Microbiology 62, 2565–2571.

Slobodkin AI, Reysenbach A-L, Slobodkina GB, Kolganova TV, Kostrikina NA, Bonch-Osmolovskaya EA (2013) Dissulfuribacter thermophilus gen. nov., sp. nov., a thermophilic, autotrophic, sulfur-disproportionating, deeply branching deltaproteobacterium from a deep-sea hydrothermal vent. International Journal of Systematic and Evolutionary Microbiology 63, 1967–1971.

Slobodkin AI, Slobodkina GB (2019) Diversity of Sulfur-Disproportionating Microorganisms. Microbiology 88, 509–522.

Slobodkina GB, Kolganova TV, Kopitsyn DS, Viryasov MB, Bonch-Osmolovskaya EA, Slobodkin AI (2016) Dissulfurirhabdus thermomarina gen. nov., sp. nov., a thermophilic, autotrophic, sulfite-reducing and disproportionating deltaproteobacterium isolated from a shallow-sea hydrothermal vent. International Journal of Systematic and Evolutionary Microbiology 66, 2515–2519.

Slobodkina GB, Reysenbach AL, Kolganova TV, Novikov AA, Bonch-Osmolovskaya EA, Slobodkin AI (2017) Thermosulfuriphilus ammonigenes gen. Nov., sp. nov., a thermophilic, chemolithoautotrophic bacterium capable of respiratory ammonification of nitrate with elemental sulfur. International Journal of Systematic and Evolutionary Microbiology 67, 3474–3479.

Sorokin DYu, Tourova TP, Mußmann M, Muyzer G (2008) Dethiobacter alkaliphilus gen. nov. sp. nov., and Desulfurivibrio alkaliphilus gen. nov. sp. nov.: two novel representatives of reductive sulfur cycle from soda lakes. Extremophiles 12, 431–439.

Stookey LL (1970) Ferrozine-A New Spectrophotometric Reagent for Iron. Analytical Chemistry 42, 779–781.

Thamdrup B, Finster K, Hansen JW, Bak F (1993) Bacterial disproportionation of elemental sulfur coupled to chemical reduction of iron or manganese. Applied and Environmental Microbiology 59, 101–108.

Umezawa K, Kojima H, Kato Y, Fukui M (2021) Dissulfurispira thermophila gen. nov., sp. nov., a thermophilic chemolithoautotroph growing by sulfur disproportionation, and proposal of novel taxa in the phylum Nitrospirota to reclassify the genus Thermodesulfovibrio. Systematic and Applied Microbiology 44.

Varghese NJ, Mukherjee S, Ivanova N, Konstantinidis KT, Mavrommatis K, Kyrpides NC, Pati A (2015) Microbial species delineation using whole genome sequences. Nucleic Acids Research 43, 6761–6771.

Waite DW, Chuvochina M, Pelikan C, Parks DH, Yilmaz P, Wagner M, Loy A, Naganuma T, Nakai R, Whitman WB, Hahn MW, Kuever J, Hugenholtz P (2020) Proposal to reclassify the proteobacterial classes Deltaproteobacteria and Oligoflexia, and the phylum Thermodesulfobacteria into four phyla reflecting major functional capabilities. International Journal of Systematic and Evolutionary Microbiology 70, 5972–6016.

Ward LM, Bertran E, Johnston DT (2021) Expanded Genomic Sampling Refines Current Understanding of the Distribution and Evolution of Sulfur Metabolisms in the Desulfobulbales. Frontiers in Microbiology 12.

Widdel F, Pfennig N (1982) Studies on dissimilatory sulfate-reducing bacteria that decompose fatty acids II. Incomplete oxidation of propionate by Desulfobulbus propionicus gen. nov., sp. nov. Archives of Microbiology 131, 360–365.

Yarza P, Yilmaz P, Pruesse E, Glöckner FO, Ludwig W, Schleifer K-H, Whitman WB, Euzéby J, Amann R, Rosselló-Móra R (2014) Uniting the classification of cultured and uncultured bacteria and archaea using 16S rRNA gene sequences. Nature Reviews Microbiology 12, 635–645.

Zhou Z, Tran P, Liu Y, Kieft K, Anantharaman K (2019) METABOLIC: A scalable high-throughput metabolic and biogeochemical functional trait profiler based on microbial genomes.

