## Supplemental File 1 for "*Thiovibrio frasassiensis* gen. nov., sp. nov., an autotrophic, elemental sulfur disproportionating bacterium isolated from sulfidic karst sediment, and proposal of Thiovibrionaceae fam. nov."

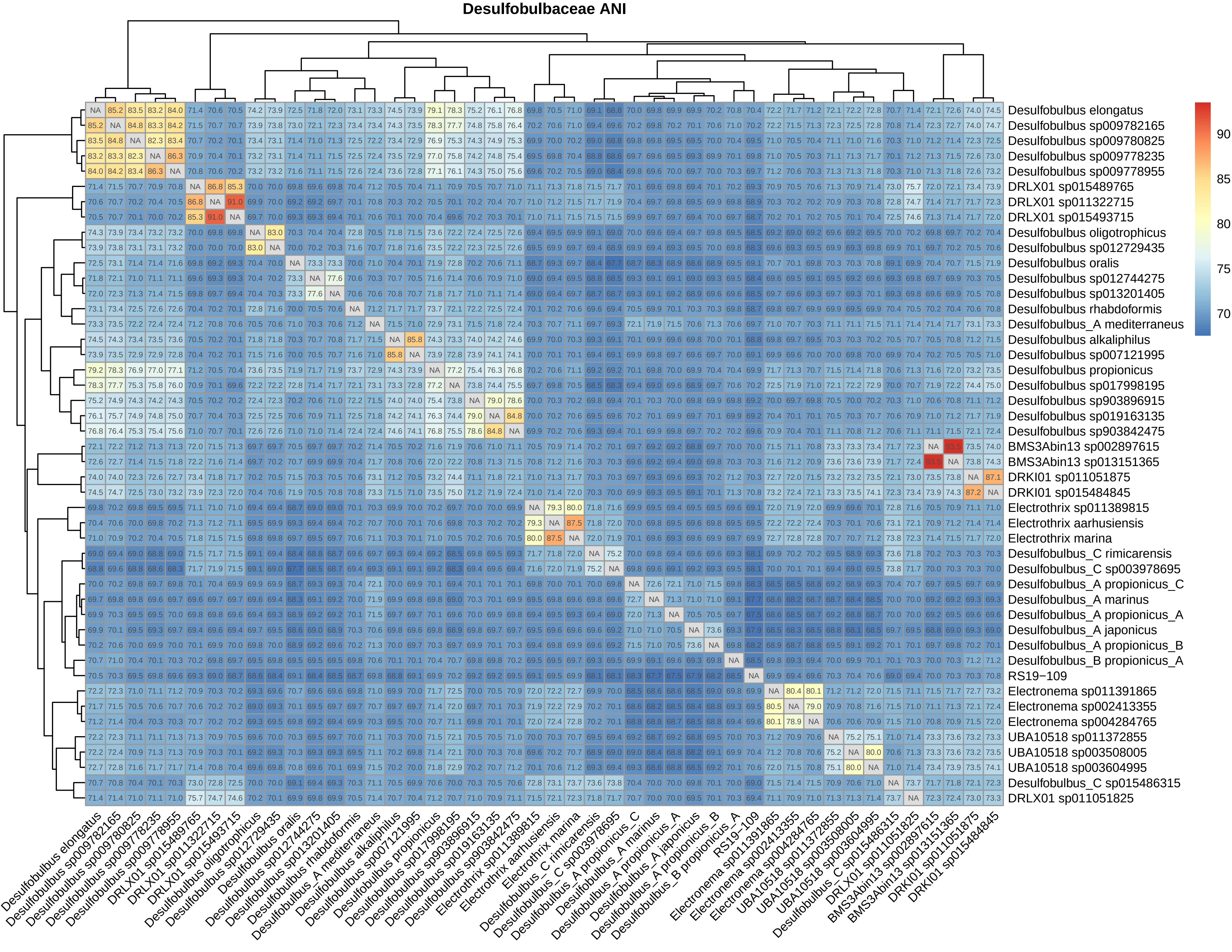
Desulfobulbaceae ANI
90
85
80
75
70
Desulfobulbus elongatus
Desulfobulbus sp009782165
Desulfobulbus sp009780825
Desulfobulbus sp009778235
Desulfobulbus sp009778955
DRLX01 sp015489765
DRLX01 sp011322715
DRLX01 sp015493715
Desulfobulbus oligotrophicus
Desulfobulbus sp012729435
Desulfobulbus oralis
Desulfobulbus sp012744275
Desulfobulbus sp013201405
Desulfobulbus rhabdoformis
Desulfobulbus_A mediterraneus
Desulfobulbus alkaliphilus
Desulfobulbus sp007121995
Desulfobulbus propionicus
Desulfobulbus sp017998195
Desulfobulbus sp903896915
Desulfobulbus sp019163135
Desulfobulbus sp903842475
BMS3Abin13 sp002897615
BMS3Abin13 sp013151365
DRKI01 sp011051875
DRKI01 sp015484845
Electrothrix sp011389815
Electrothrix aarhusiensis
Electrothrix marina
Desulfobulbus_C rimicarensis
Desulfobulbus_C sp003978695
Desulfobulbus_A propionicus_C
Desulfobulbus_A marinus
Desulfobulbus_A propionicus_A
Desulfobulbus_A japonicus
Desulfobulbus_A propionicus_B
Desulfobulbus_B propionicus_A
RS19−109
Electronema sp011391865
Electronema sp002413355
Electronema sp004284765
UBA10518 sp011372855
UBA10518 sp003508005
UBA10518 sp003604995
Desulfobulbus_C sp015486315
DRLX01 sp011051825

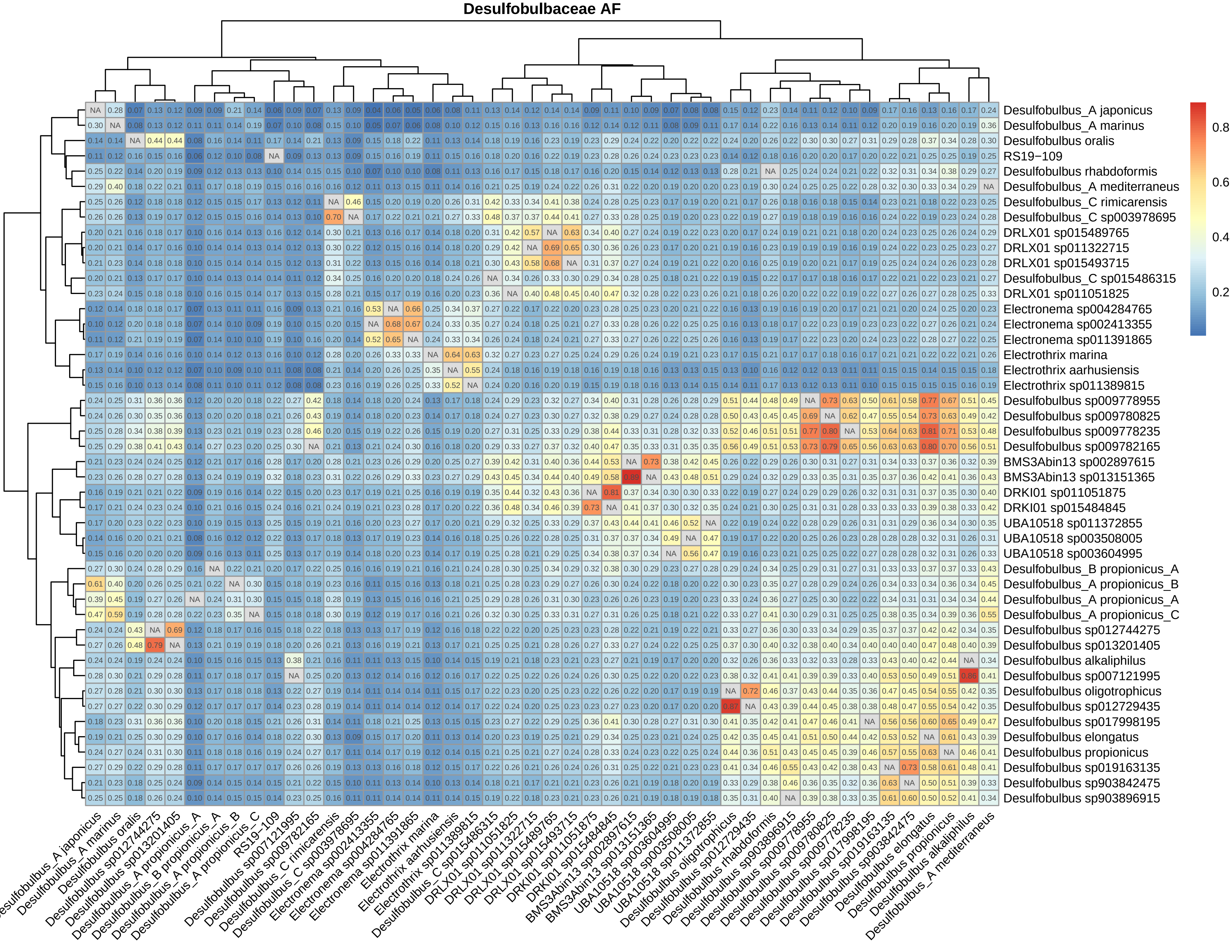

Desulfobulbaceae AF
0.8
0.6
0.4
0.2
Desulfobulbus_A japonicus
Desulfobulbus_A marinus
Desulfobulbus oralis
RS19−109
Desulfobulbus rhabdoformis
Desulfobulbus_A mediterraneus
Desulfobulbus_C rimicarensis
Desulfobulbus_C sp003978695
DRLX01 sp015489765
DRLX01 sp011322715
DRLX01 sp015493715
Desulfobulbus_C sp015486315
DRLX01 sp011051825
Electronema sp004284765
Electronema sp002413355
Electronema sp011391865
Electrothrix marina
Electrothrix aarhusiensis
Electrothrix sp011389815
Desulfobulbus sp009778955
Desulfobulbus sp009780825
Desulfobulbus sp009778235
Desulfobulbus sp009782165
BMS3Abin13 sp002897615
BMS3Abin13 sp013151365
DRKI01 sp011051875
DRKI01 sp015484845
UBA10518 sp011372855
UBA10518 sp003508005
UBA10518 sp003604995
Desulfobulbus_B propionicus_A
Desulfobulbus_A propionicus_B
Desulfobulbus_A propionicus_A
Desulfobulbus_A propionicus_C
Desulfobulbus sp012744275
Desulfobulbus sp013201405
Desulfobulbus alkaliphilus
Desulfobulbus sp007121995
Desulfobulbus oligotrophicus
Desulfobulbus sp012729435
Desulfobulbus sp017998195
Desulfobulbus elongatus
Desulfobulbus propionicus
Desulfobulbus sp019163135
Desulfobulbus sp903842475
Desulfobulbus sp903896915

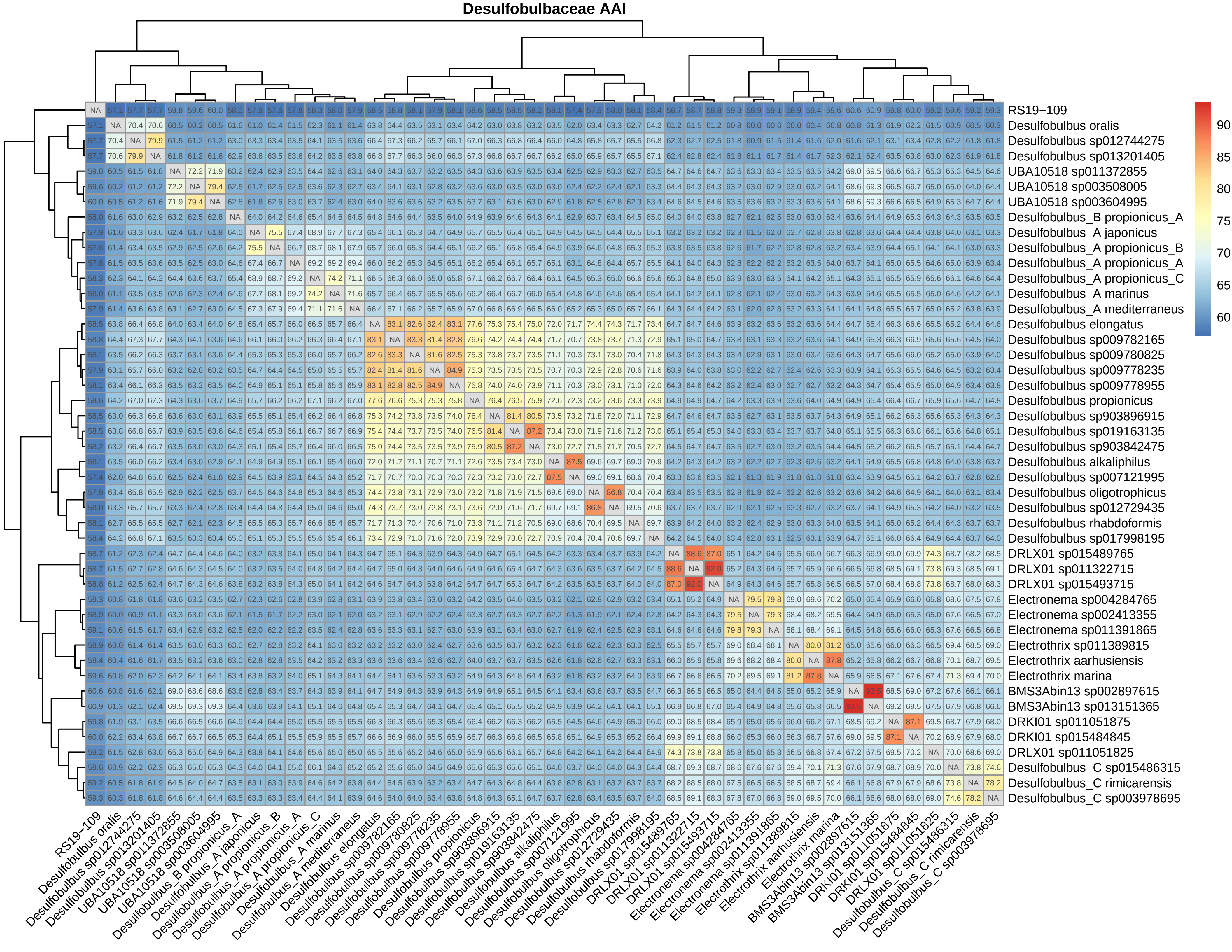
Desulfobulbaceae AAI
RS19−109
Desulfobulbus oralis
Desulfobulbus sp012744275
Desulfobulbus sp013201405
UBA10518 sp011372855
UBA10518 sp003508005
UBA10518 sp003604995
Desulfobulbus_B propionicus_A
Desulfobulbus_A japonicus
Desulfobulbus_A propionicus_B
Desulfobulbus_A propionicus_A
Desulfobulbus_A propionicus_C
Desulfobulbus_A marinus
Desulfobulbus_A mediterraneus
Desulfobulbus elongatus
Desulfobulbus sp009782165
Desulfobulbus sp009780825
Desulfobulbus sp009778235
Desulfobulbus sp009778955
Desulfobulbus propionicus
Desulfobulbus sp903896915
Desulfobulbus sp019163135
Desulfobulbus sp903842475
Desulfobulbus alkaliphilus
Desulfobulbus sp007121995
Desulfobulbus oligotrophicus
Desulfobulbus sp012729435
Desulfobulbus rhabdoformis
Desulfobulbus sp017998195
DRLX01 sp015489765
DRLX01 sp011322715
DRLX01 sp015493715
Electronema sp004284765
Electronema sp002413355
Electronema sp011391865
Electrothrix sp011389815
Electrothrix aarhusiensis
Electrothrix marina
BMS3Abin13 sp002897615
BMS3Abin13 sp013151365
DRKI01 sp011051875
DRKI01 sp015484845
DRLX01 sp011051825
Desulfobulbus_C sp015486315
Desulfobulbus_C rimicarensis
Desulfobulbus_C sp003978695
90
85
80
75
70
65
60

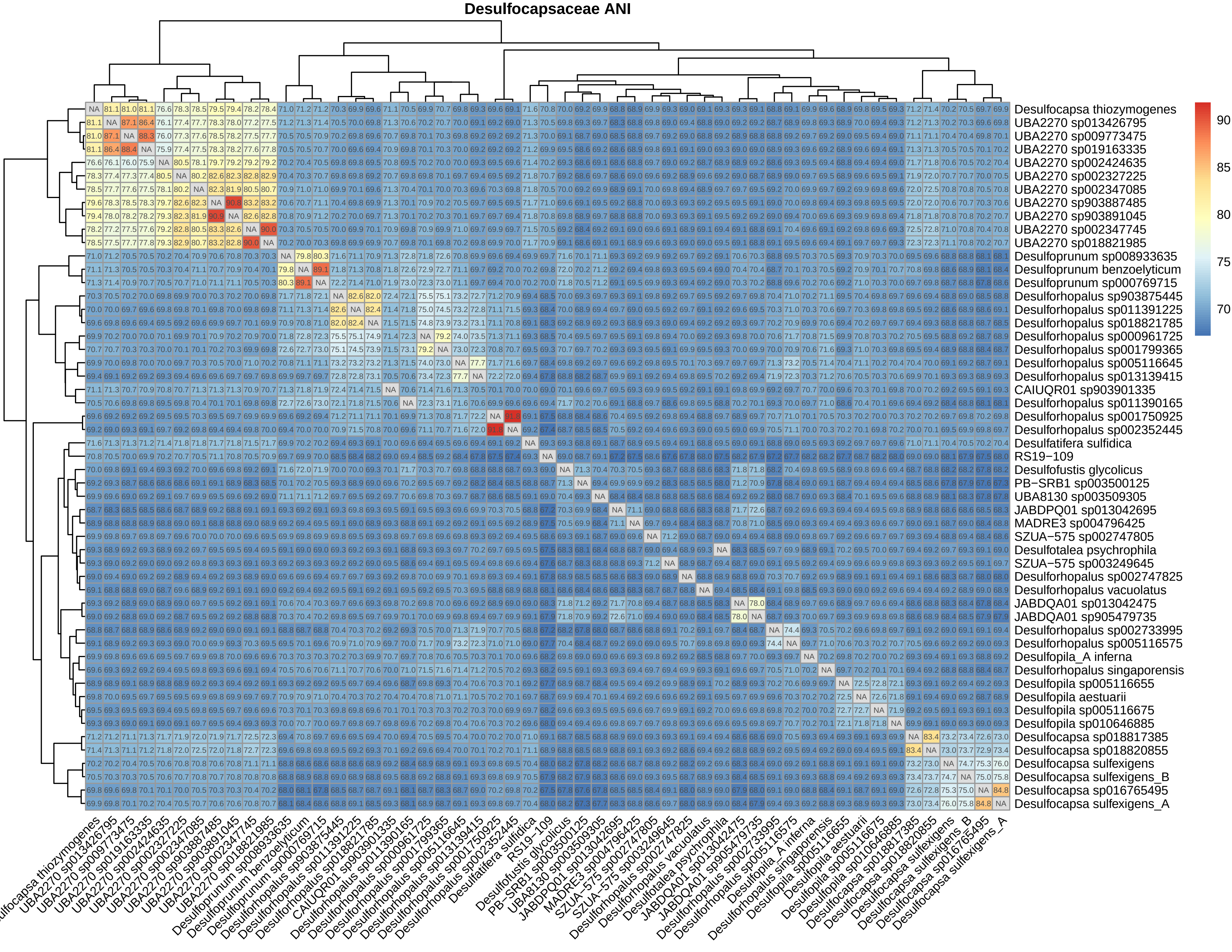

Desulfocapsaceae ANI
90
85
80
75
70
Desulfocapsa thiozymogenes
UBA2270 sp013426795
UBA2270 sp009773475
UBA2270 sp019163335
UBA2270 sp002424635
UBA2270 sp002327225
UBA2270 sp002347085
UBA2270 sp903887485
UBA2270 sp903891045
UBA2270 sp002347745
UBA2270 sp018821985
Desulfoprunum sp008933635
Desulfoprunum benzoelyticum
Desulfoprunum sp000769715
Desulforhopalus sp903875445
Desulforhopalus sp011391225
Desulforhopalus sp018821785
Desulforhopalus sp000961725
Desulforhopalus sp001799365
Desulforhopalus sp005116645
Desulforhopalus sp013139415
CAIUQR01 sp903901335
Desulforhopalus sp011390165
Desulforhopalus sp001750925
Desulforhopalus sp002352445
Desulfatifera sulfidica
RS19−109
Desulfofustis glycolicus
PB−SRB1 sp003500125
UBA8130 sp003509305
JABDPQ01 sp013042695
MADRE3 sp004796425
SZUA−575 sp002747805
Desulfotalea psychrophila
SZUA−575 sp003249645
Desulforhopalus sp002747825
Desulforhopalus vacuolatus
JABDQA01 sp013042475
JABDQA01 sp905479735
Desulforhopalus sp002733995
Desulforhopalus sp005116575
Desulfopila_A inferna
Desulforhopalus singaporensis
Desulfopila sp005116655
Desulfopila aestuarii
Desulfopila sp005116675
Desulfopila sp010646885
Desulfocapsa sp018817385
Desulfocapsa sp018820855
Desulfocapsa sulfexigens
Desulfocapsa sulfexigens_B
Desulfocapsa sp016765495
Desulfocapsa sulfexigens_A

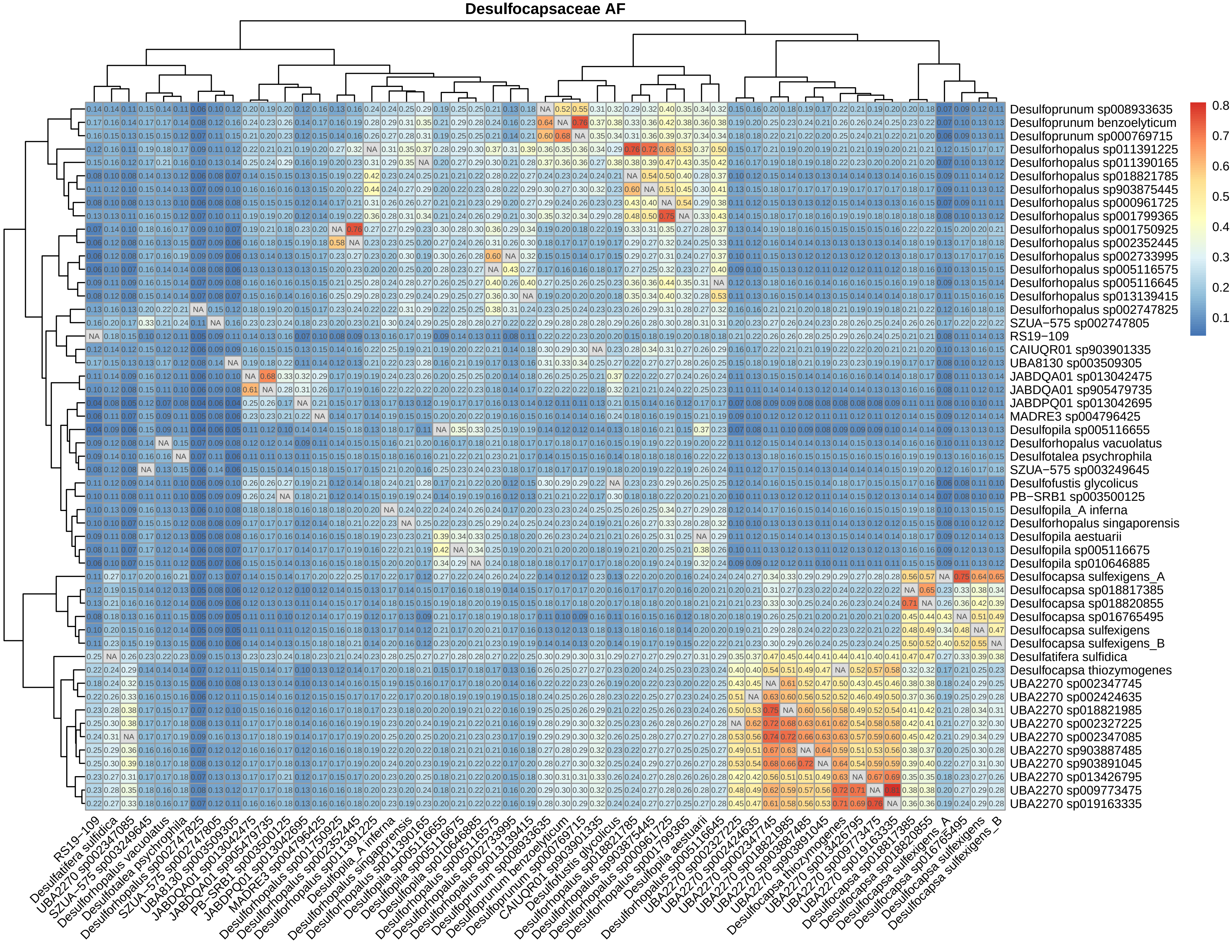
Desulfocapsaceae AF
Desulfoprunum sp008933635
Desulfoprunum benzoelyticum
Desulfoprunum sp000769715
Desulforhopalus sp011391225
Desulforhopalus sp011390165
Desulforhopalus sp018821785
Desulforhopalus sp903875445
Desulforhopalus sp000961725
Desulforhopalus sp001799365
Desulforhopalus sp001750925
Desulforhopalus sp002352445
Desulforhopalus sp002733995
Desulforhopalus sp005116575
Desulforhopalus sp005116645
Desulforhopalus sp013139415
Desulforhopalus sp002747825
SZUA−575 sp002747805
RS19−109
CAIUQR01 sp903901335
UBA8130 sp003509305
JABDQA01 sp013042475
JABDQA01 sp905479735
JABDPQ01 sp013042695
MADRE3 sp004796425
Desulfopila sp005116655
Desulforhopalus vacuolatus
Desulfotalea psychrophila
SZUA−575 sp003249645
Desulfofustis glycolicus
PB−SRB1 sp003500125
Desulfopila_A inferna
Desulforhopalus singaporensis
Desulfopila aestuarii
Desulfopila sp005116675
Desulfopila sp010646885
Desulfocapsa sulfexigens_A
Desulfocapsa sp018817385
Desulfocapsa sp018820855
Desulfocapsa sp016765495
Desulfocapsa sulfexigens
Desulfocapsa sulfexigens_B
Desulfatifera sulfidica
Desulfocapsa thiozymogenes
UBA2270 sp002347745
UBA2270 sp002424635
UBA2270 sp018821985
UBA2270 sp002327225
UBA2270 sp002347085
UBA2270 sp903887485
UBA2270 sp903891045
UBA2270 sp013426795
UBA2270 sp009773475
UBA2270 sp019163335
0.8
0.7
0.6
0.5
0.4
0.3
0.2
0.1

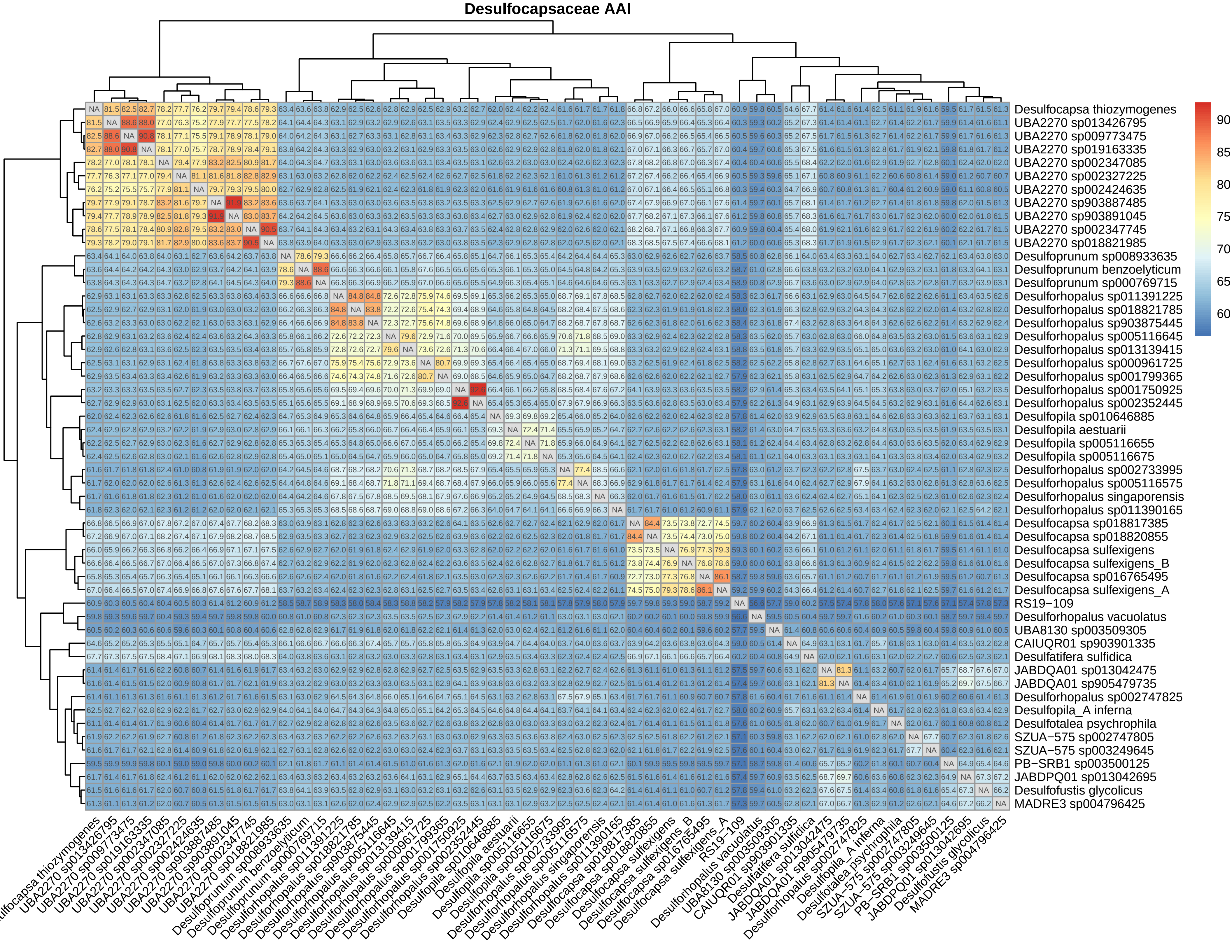

Desulfocapsaceae AAI
90
85
80
75
70
65
60
Desulfocapsa thiozymogenes
UBA2270 sp013426795
UBA2270 sp009773475
UBA2270 sp019163335
UBA2270 sp002347085
UBA2270 sp002327225
UBA2270 sp002424635
UBA2270 sp903887485
UBA2270 sp903891045
UBA2270 sp002347745
UBA2270 sp018821985
Desulfoprunum sp008933635
Desulfoprunum benzoelyticum
Desulfoprunum sp000769715
Desulforhopalus sp011391225
Desulforhopalus sp018821785
Desulforhopalus sp903875445
Desulforhopalus sp005116645
Desulforhopalus sp013139415
Desulforhopalus sp000961725
Desulforhopalus sp001799365
Desulforhopalus sp001750925
Desulforhopalus sp002352445
Desulfopila sp010646885
Desulfopila aestuarii
Desulfopila sp005116655
Desulfopila sp005116675
Desulforhopalus sp002733995
Desulforhopalus sp005116575
Desulforhopalus singaporensis
Desulforhopalus sp011390165
Desulfocapsa sp018817385
Desulfocapsa sp018820855
Desulfocapsa sulfexigens
Desulfocapsa sulfexigens_B
Desulfocapsa sp016765495
Desulfocapsa sulfexigens_A
RS19−109
Desulforhopalus vacuolatus
UBA8130 sp003509305
CAIUQR01 sp903901335
Desulfatifera sulfidica
JABDQA01 sp013042475
JABDQA01 sp905479735
Desulforhopalus sp002747825
Desulfopila_A inferna
Desulfotalea psychrophila
SZUA−575 sp002747805
SZUA−575 sp003249645
PB−SRB1 sp003500125
JABDPQ01 sp013042695
Desulfofustis glycolicus
MADRE3 sp004796425

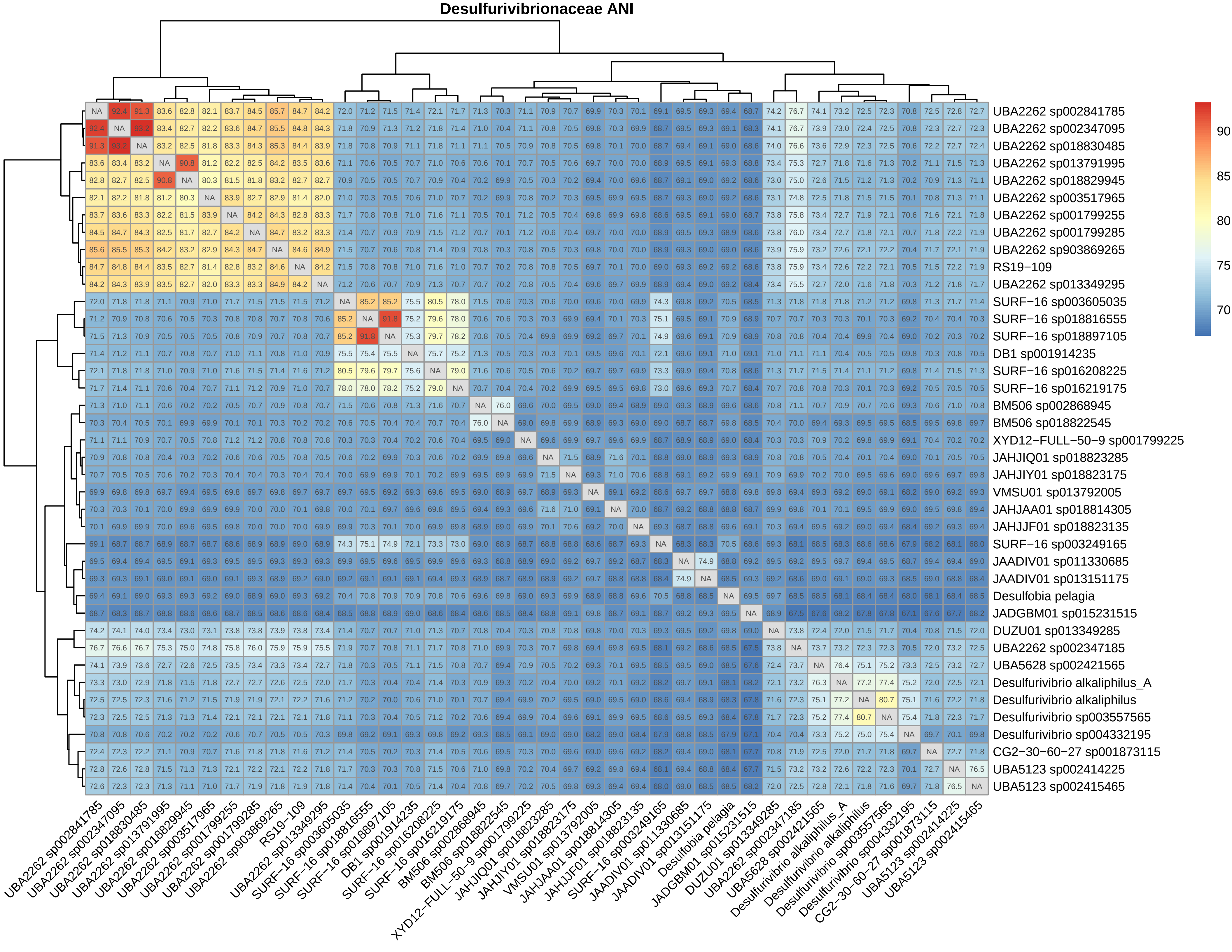
Desulfurivibrionaceae ANI
UBA2262 sp002841785
UBA2262 sp002347095
UBA2262 sp018830485
UBA2262 sp013791995
UBA2262 sp018829945
UBA2262 sp003517965
UBA2262 sp001799255
UBA2262 sp001799285
UBA2262 sp903869265
RS19−109
UBA2262 sp013349295
SURF−16 sp003605035
SURF−16 sp018816555
SURF−16 sp018897105
DB1 sp001914235
SURF−16 sp016208225
SURF−16 sp016219175
BM506 sp002868945
BM506 sp018822545
XYD12−FULL−50−9 sp001799225
JAHJIQ01 sp018823285
JAHJIY01 sp018823175
VMSU01 sp013792005
JAHJAA01 sp018814305
JAHJJF01 sp018823135
SURF−16 sp003249165
JAADIV01 sp011330685
JAADIV01 sp013151175
Desulfobia pelagia
JADGBM01 sp015231515
DUZU01 sp013349285
UBA2262 sp002347185
UBA5628 sp002421565
Desulfurivibrio alkaliphilus_A
Desulfurivibrio alkaliphilus
Desulfurivibrio sp003557565
Desulfurivibrio sp004332195
CG2−30−60−27 sp001873115
UBA5123 sp002414225
UBA5123 sp002415465
90
85
80
75
70

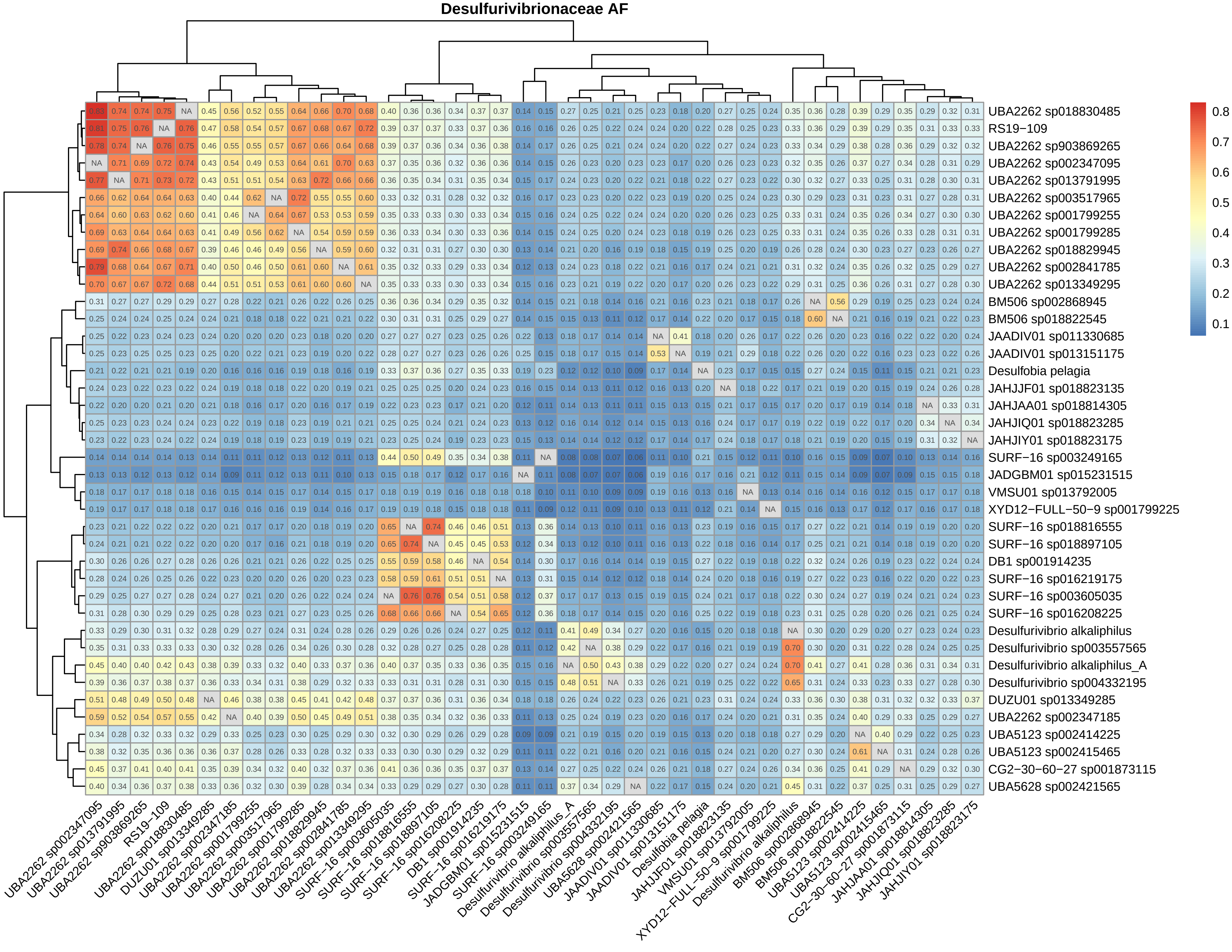
Desulfurivibrionaceae AF
UBA2262 sp018830485
RS19−109
UBA2262 sp903869265
UBA2262 sp002347095
UBA2262 sp013791995
UBA2262 sp003517965
UBA2262 sp001799255
UBA2262 sp001799285
UBA2262 sp018829945
UBA2262 sp002841785
UBA2262 sp013349295
BM506 sp002868945
BM506 sp018822545
JAADIV01 sp011330685
JAADIV01 sp013151175
Desulfobia pelagia
JAHJJF01 sp018823135
JAHJAA01 sp018814305
JAHJIQ01 sp018823285
JAHJIY01 sp018823175
SURF−16 sp003249165
JADGBM01 sp015231515
VMSU01 sp013792005
XYD12−FULL−50−9 sp001799225
SURF−16 sp018816555
SURF−16 sp018897105
DB1 sp001914235
SURF−16 sp016219175
SURF−16 sp003605035
SURF−16 sp016208225
Desulfurivibrio alkaliphilus
Desulfurivibrio sp003557565
Desulfurivibrio alkaliphilus_A
Desulfurivibrio sp004332195
DUZU01 sp013349285
UBA2262 sp002347185
UBA5123 sp002414225
UBA5123 sp002415465
CG2−30−60−27 sp001873115
UBA5628 sp002421565
0.8
0.7
0.6
0.5
0.4
0.3
0.2
0.1

**Desulfurivibrionaceae AAI**

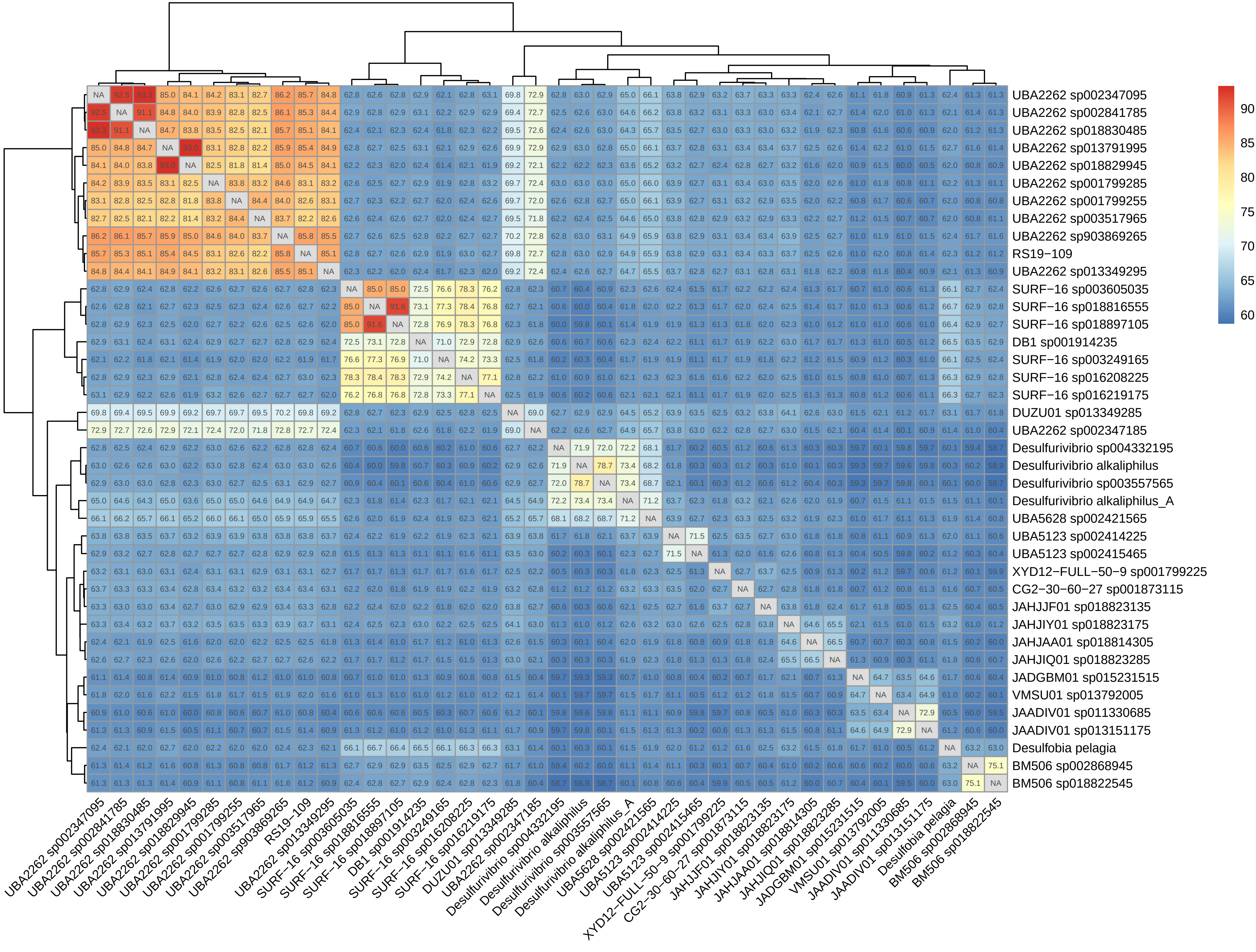

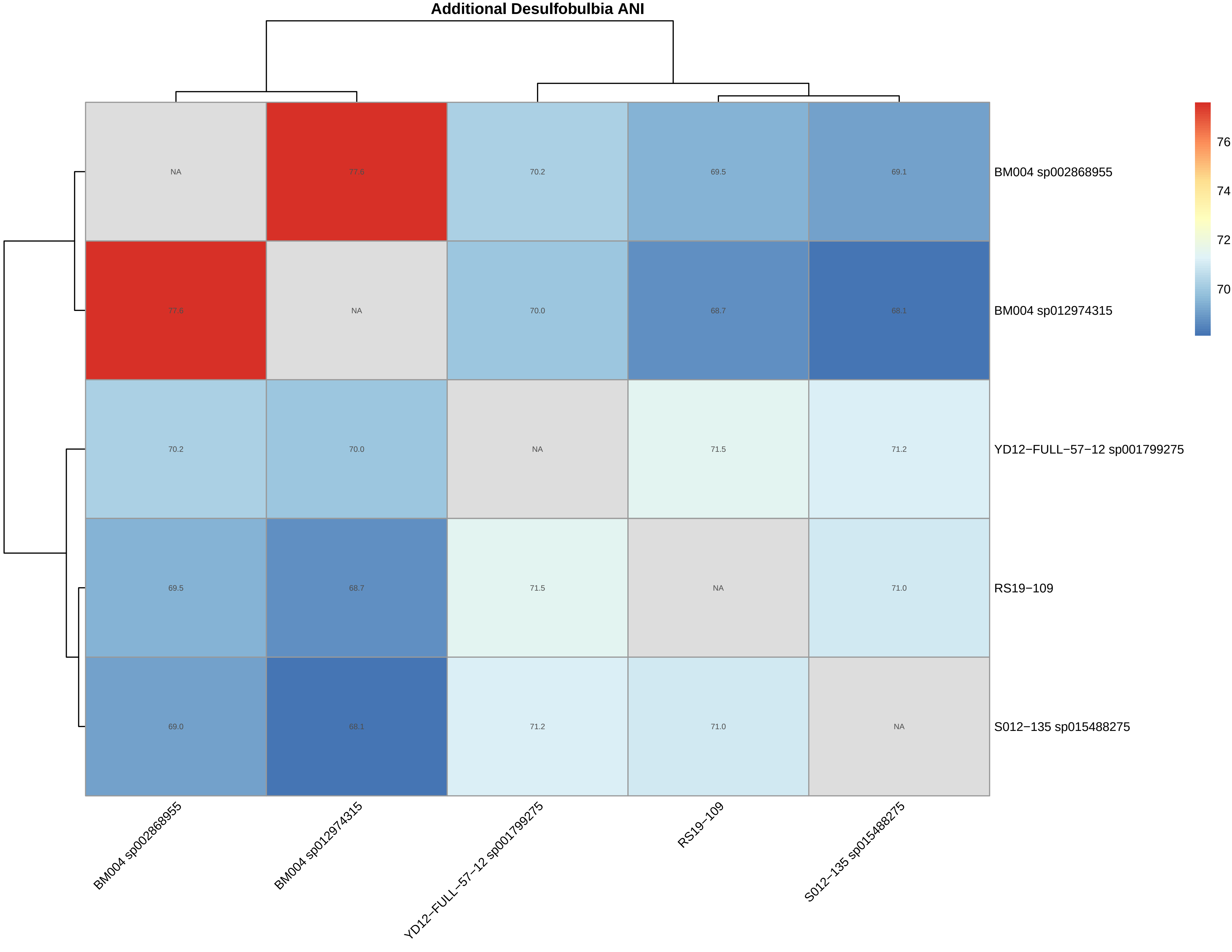
Additional Desulfobulbia ANI
BM004 sp002868955
BM004 sp012974315
YD12−FULL−57−12 sp001799275
RS19−109
S012−135 sp015488275
NA
77.6
70.2
69.5
69.1
77.6
NA
70.0
68.7
68.1
70.2
70.0
NA
71.5
71.2
69.5
68.7
71.5
NA
71.0
69.0
68.1
71.2
71.0
NA
76
74
72
70

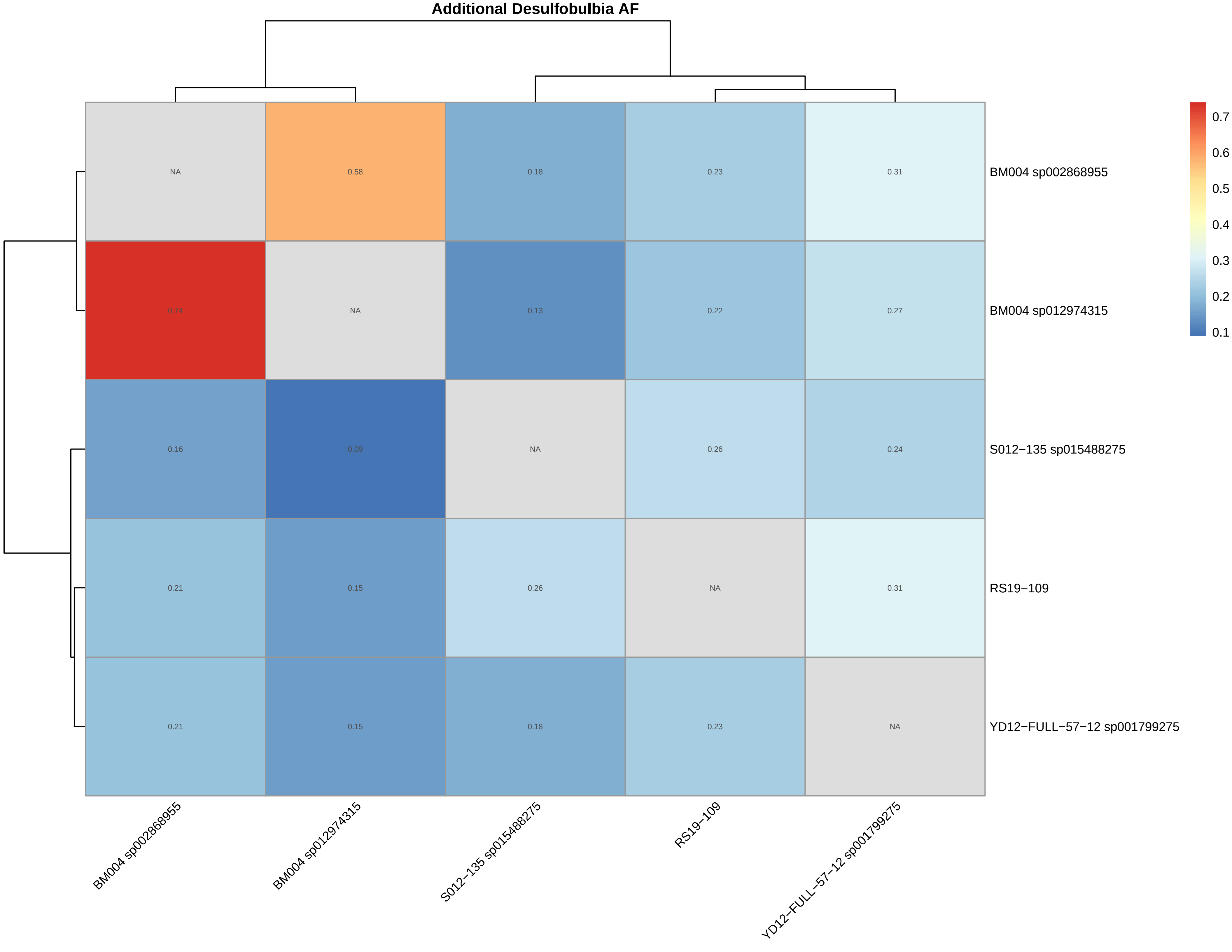

Additional Desulfobulbia AF
BM004 sp002868955
BM004 sp012974315
S012−135 sp015488275
RS19−109
YD12−FULL−57−12 sp001799275
NA
0.58
0.18
0.23
0.31
0.74
NA
0.13
0.22
0.27
0.16
0.09
NA
0.26
0.24
0.21
0.15
0.26
NA
0.31
0.21
0.15
0.18
0.23
NA
0.7
0.6
0.5
0.4
0.3
0.2
0.1

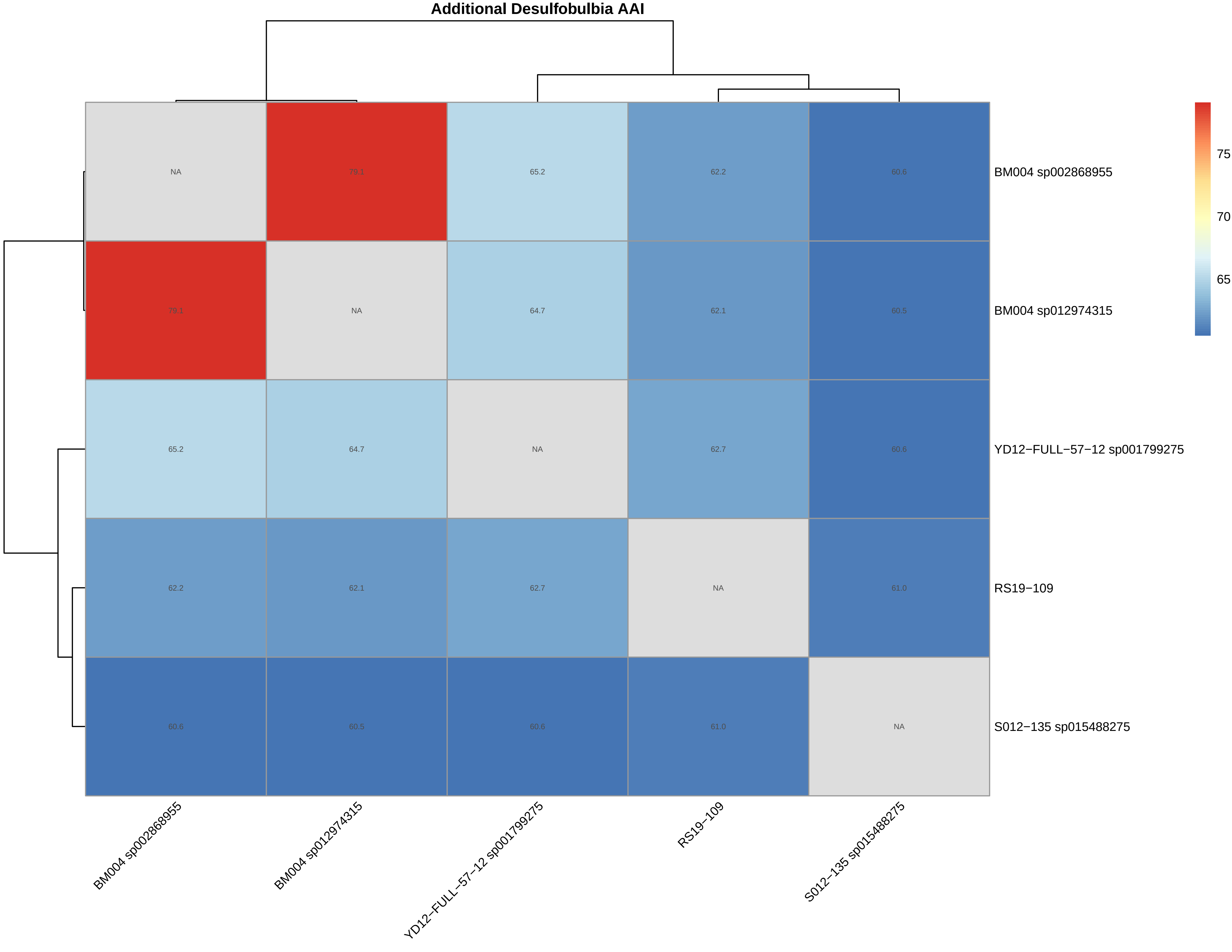
Additional Desulfobulbia AAI
BM004 sp002868955
BM004 sp012974315
YD12−FULL−57−12 sp001799275
RS19−109
S012−135 sp015488275
NA
79.1
65.2
62.2
60.6
64.7
62.1
60.5
62.7
61.0
75
70
65
