## Supplementary figures and images for "*Thiovibrio frasassiensis* gen. nov., sp. nov., an autotrophic, elemental sulfur disproportionating bacterium isolated from sulfidic karst sediment, and proposal of Thiovibrionaceae fam. nov."

### Supplemental File 2

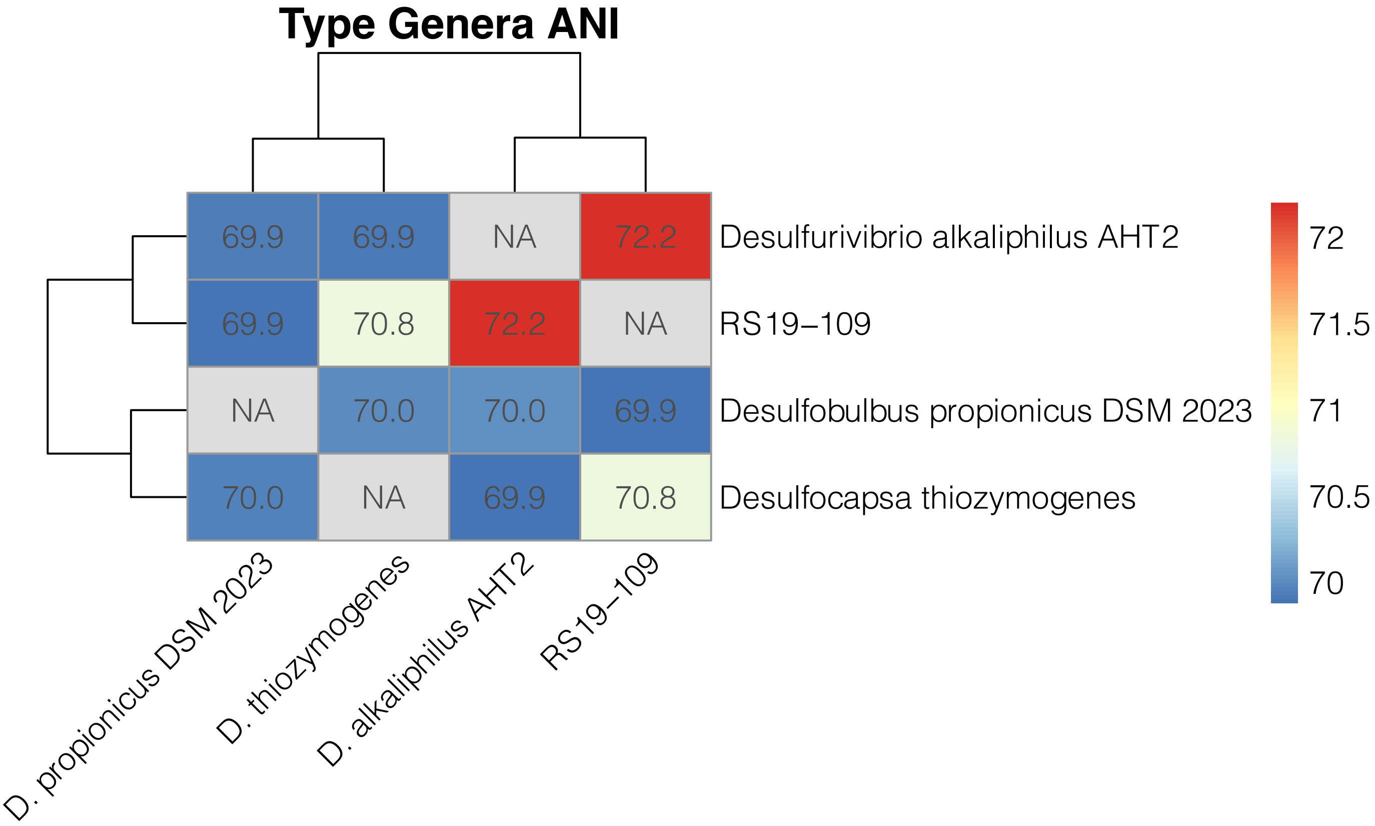

#### Type Genera AAI

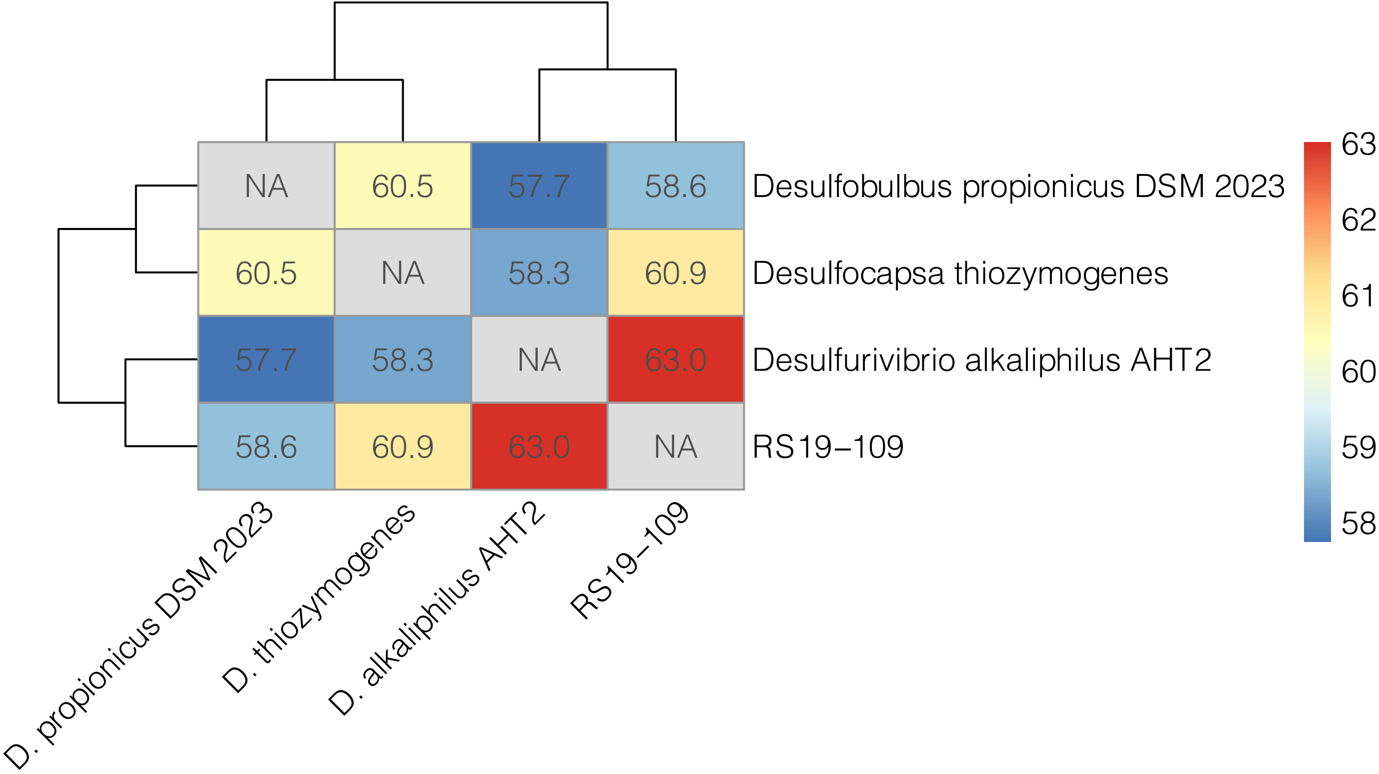
